# Bergerac Strains of *C. elegans* Revisited: Expansion of Tc*1* elements Impose a Significant Genomic and Fitness Cost

**DOI:** 10.1101/2022.02.02.478770

**Authors:** Austin T. Daigle, Thaddeus C. Deiss, Robert H. Melde, Ulfar Bergthorsson, Vaishali Katju

**Affiliations:** Department of Veterinary Integrative Biosciences, College of Veterinary Medicine and Biomedical Sciences, Texas A&M University, College Station, TX 77845, USA; Department of Integrative Biology, University of Wisconsin-Madison, Madison, WI 53706, USA

**Author notes:** Joint first authors (Austin T. Daigle, Thaddeus C. Deiss). **Data Availability:** Whole genome sequence files are available in fastq format from the NCBI SRA database under Bioproject PRJNA801693 (https://www.ncbi.nlm.nih.gov/sra/PRJNA801693). Accessions for individual strains sequenced in this study are as follows: CB4851 (SAMN25375976), RW6999 (SAMN25375977), RW7000 (SAMN25375978) and Bristol N2 (SAMN25375979).

**Keywords:** *Caenorhabditis elegans*, Bergerac, transposable element, transposon, copy-number variation, fitness, whole-genome sequencing, chromatin, recombination

## Abstract

The DNA transposon Tc*1* was the first transposable element (TE) to be characterized in *Caenorhabditis elegans* and to date, remains the best-studied TE in *Caenorhabditis* worms. While Tc*1* copy-number is regulated at approximately 30 copies in the laboratory N2/Bristol and the vast majority of *C. elegans* strains, the Bergerac strain and its derivatives have experienced a marked Tc*1* proliferation. Given the historical importance of the Bergerac strain in the development of the *C. elegans* model, we implemented a modern genomic analysis of three Bergerac strains (CB4851, RW6999, and RW7000) in conjunction with multiple phenotypic assays to better elucidate the (i) genomic distribution of Tc*1*, and (ii) phenotypic consequences of TE deregulation for the host organism. The median estimates of Tc*1* copy-number in the Bergerac strains ranged from 451 to 748, which is both (i) greater than previously estimated, and (ii) likely to be an underestimate of the actual copy-numbers since coverage-based estimates and ddPCR results both suggest higher Tc*1* numbers. All three Bergerac strains had significantly reduced trait means compared to the N2 control for each of four fitness-related traits, with specific traits displaying significant differences between Bergerac strains. T*c1* proliferation was genome-wide, specific to Tc*1*, and particularly high on chromosomes V and X. There were fewer Tc*1* insertions in highly expressed chromatin environments than expected by chance. Furthermore, Tc*1* integration motifs were also less frequent in exon than non-coding sequences. The source of the proliferation of Tc*1* in the Bergerac strains is specific to Tc*1* and independent of other TEs. The Bergerac strains contain none of the alleles that have previously been found to derepress TE activity in *C. elegans*. However, the Bergerac strains had several Tc*1* insertions near or within highly germline-transcribed genes which could account for the recent germline proliferation.

## INTRODUCTION

Transposable elements (TEs henceforth) are small DNA fragments particularly ubiquitous in eukaryotic genomes that occasionally relocate to new genomic position, hence the moniker of “jumping genes” (McClintock 1948; Chuong *et al*. 2016). TEs are viewed as parasitic and selfish because they rely on the host genome’s cellular machinery to mobilize, and their ability to spread despite deleterious fitness costs associated with disruption of host gene function due to insertions (Orgel and Crick 1980; Muñoz-López and García-Pérez 2010). As such, both prokaryotic and eukaryotic host genomes have ubiquitously evolved complex surveillance and regulatory systems for TE detection and silencing (Sijen and Plasterk 2003; Blumenstiel 2011; Klein and O’Neill 2018; Payer and Burns 2019). Despite their overwhelmingly deleterious effects for host genomes, these mobile genetic elements can also inadvertently contribute to novel advantageous phenotypes as documented in the rapid phenotypic evolution observed in the British peppered moth (van’t Hof *et al*. 2011), the role of TEs in gene transcription as *cis*-regulatory elements in many organisms (Rebollo *et al*. 2012) and their domestication as a host recombinase mechanism required for genomic rearrangements integral to the adaptive immune systems of higher vertebrate (Yuhang *et al*. 2019). Recent research has utilized increasingly sophisticated computational tools and a comparative genomics framework to understand the broad genomic effects and functions of TEs (Bourque *et al*. 2018) while emphasizing the need for studies focusing on the initial proliferation of diverse TE families in diverse taxa (Wells and Feschotte 2020). To fully understand the coevolution between TE regulation and expansion in evolutionary diversification, the mechanisms by which TEs evade regulation at the expense of the host organism requires further elucidation.

The nematode *Caenorhabditis elegans* has been a useful model for TE research since the discovery of the first *C. elegans* TE, a DNA transposon named Tc*1* (Emmons *et al*. 1983). Tc*1* and its homologs are among the most abundant TEs occurring in nature (Plasterk *et al*. 1999). Overall, TEs comprise approximately 12% of the *C. elegans* genome (*C. elegans* Sequencing Consortium 1998). A recent study of TEs in the genomes of 208 *C. elegans* wild isolates revealed significant variation in the copy-number of diverse TE classes (Laricchia *et al*. 2017), highlighting the contribution of TEs to intraspecific genome variation. Transcription of TEs is under strong purifying selection in *C. elegans* (Bergthorsson *et al*. 2020). The primary mechanisms responsible for the regulation of transposons and other foreign DNA in the *C. elegans* genome are composed of small RNAs (sRNAs), including 21U-RNAs, 26G-RNAs and 22G-RNAs (Ambros *et al*. 2003), that couple with and guide Argonaute (AGO) proteins (Yigit *et al*. 2006).

Nematode sRNAs have distinct structures and biogenesis pathways, but often share common regulatory functions (Almeida *et al*. 2019). 21U-RNAs, a special class of sRNA known as PIWI-interacting or piRNAs, were determined to interact directly with the AGO protein *prg-1* to prevent TE spread in germ line cells (Batista *et al*. 2008; Bagijn *et al*. 2012). Despite evidence that 21U-RNAs play a role in controlling TE transcription, only the mariner transposon *Tc3* has been demonstrated to become mobilized upon disruption of piRNA machinery (Das *et al*. 2008; Reed *et al*. 2020). High-throughput sequencing of piRNAs and siRNAs in various mutants seemed to suggest that WAGO-class 22G-RNAs played a larger role in transposon silencing, and it had been suggested that the mutants studied could have inherited long-term effects of piRNA silencing due to multigenerational epigenetic memories (Ashe *et al*. 2012; Reed *et al*. 2020). However, a recent experimental study in *C. elegans* demonstrated that 22G-RNA epimutations are typically unstable with an average persistence time of two to three generations (Beltran *et al*. 2020). Many details and overlapping features of the intricate regulatory pathways involved in silencing TEs and other germline transcripts in *C. elegans* await further elucidation.

The Bergerac strain of *C. elegans* is of historical importance in the development of the species as a model organism (Riddle *et al*. 1997). Bergerac was one of the earliest laboratory strains of *C. elegans* to be cultured and was isolated from garden soil in 1944 in Bergerac, France by Victor Nigon of the Université de Lyon (Nigon 1949; Nigon and Félix 2017). Some of the earliest genetic work in *C. elegans* occurred with the use of the Bergerac strain, including the discovery of a Mendelian temperature-sensitive allele (Fatt and Dougherty 1963). Owing to difficulties with propagation later determined to be due to temperature-sensitivity of hermaphrodites (Fatt and Dougherty 1963), infertile males (Fatt and Dougherty 1963), and high mutation rates (Moerman and Waterston 1984), the use of the Bergerac strain for subsequent genetic work was abandoned and replaced by the Bristol N2 strain isolated by L. N. Staniland from a mushroom compost heap in Bristol, England (Nicholas *et al*. 1959; Brenner 1974; Riddle *et al*. 1997; Nigon and Félix 2017). Several decades later and with a more developed arsenal of molecular and genetic tools available, several *C. elegans* researchers took to investigating the mechanistic bases of the aberrant phenotypes associated with the Bergerac strain. Bristol N2 and Bergerac were observed to exhibit differences in restriction endonuclease cleavage patterns on Southern hybridizations when probed with randomly selected cloned fragments (Emmons *et al*. 1979, 1983; Files *et al*. 1983). Southern hybridization analyses further demonstrated that while Bristol N2 possessed 20 ± 5 dispersed copies of Tc*1* in the genome, the Bergerac strain appeared to have an estimated 200 ± 50 copies (Emmons *et al*. 1983).

Further investigation of Tc*1* copy-number in ten newly available *C. elegans* isolates led Emmons *et al*. (1983) to the parsimonious conclusion that these intraspecific differences in transposon copy-number were owing to the uniquely massive proliferation of Tc*1* elements in the Bergerac strain which likely resulted in gene disruption leading to phenotypic defects. A later study using quantitative dot blot hybridization presented estimates of Tc*1* copy-number in the Bristol N2 and Bergerac strain as being ∼ 30 Tc*1* and 300-550 copies, respectively (Egilmez *et al*. 1995). Rosenzweig *et al*. (1983a) were the first to generate the complete 1,610 bp nucleotide sequence of the Tc*1* transposable element in *C. elegans* with its characteristic short inverted terminal repeats of <100 bp. Liao *et al*. (1983) provided a substantially detailed characterization of this *C. elegans* Tc*1*, namely that (i) while it shared many structural features with other eukaryotic DNA transposable elements, it belonged to a unique class, and (ii) all *C. elegans* Tc*1* copies appeared to display full-length conservation unlike *Drosophila P*-elements, suggesting that it encodes products mediating its own transcription. Tc*1* elements in the Bergerac strain were found to be significantly more active relative to their counterparts in the Bristol N2 genome, displaying site-specific insertion and excision from the muscle gene *unc-54* (Eide and Anderson 1985, 1988). In addition to increased transposition, it was noted that Bergerac strains were less fit, produced fewer progeny, moved with less coordination, and had a higher incidence of males relative to other *C. elegans* strains, despite the males being sterile (Hodgkin and Doniach 1997). It has been hypothesized that the increase in Tc*1* copy-number occurred in the laboratory after Nigon isolated Bergerac from the wild (Moerman and Waterston 1984; Egilmez *et al*. 1995; Nigon and Félix 2017). However, the cause of the massive proliferation of Tc*1* elements in the Bergerac strains remains to be identified.

While some traits in Bergerac have been studied, mainly in the strain RW7000 (Fatt and Dougherty 1963; Shook and Johnson 1999; Vertino *et al*. 2011; Lee *et al*. 2016), a comparative study of multiple fitness-related traits and precise quantification of composite fitness has yet to be conducted on multiple distinct Bergerac strains simultaneously. This project is the first to employ high-throughput Illumina whole-genome sequencing technology to sequence and analyze the entire genomes of three distinct Bergerac strains. While the genome of one Bergerac strain, CB4851, has previously been sequenced, it has not yet been analyzed to study Tc*1* proliferation (Cook *et al*. 2016; Laricchia *et al*. 2017). Herein, we quantified and compared four fitness-related traits (developmental rate, productivity, longevity, and survivorship) in order to discern extant phenotypic variation among the Bergerac strains RW6999, RW7000, and CB4851. Next, using whole-genome sequencing and ddPCR data for each strain, we generated more accurate TE copy-number estimates, quantify the decreased and variable fitness across Bergerac strains relative to Bristol N2, analyze the distribution and sequence context of Tc*1* landing sites, and conduct an initial search for the cause of increased Tc*1* proliferation.

## MATERIALS AND METHODS

### Bergerac strains of C. elegans used in this study

This study focused on three subclones of the original Bergerac strain isolated by Nigon in Bergerac, France in 1944 (Nigon and Félix 2017). This strain was shared with several other laboratories before the implementation of cryopreservation techniques for *C. elegans*, and hence the names of these subclones of the original Bergerac strain either represent their initial culture location or standardized strain nomenclature that came to be adapted later. We focused on three Bergerac strains to quantify their fitness declines and genomic divergence that may have occurred during laboratory evolution and divergence following the proliferation of Tc*1* elements. The first, RW7000 (also known as Bergerac BO for its location in Boulder, Colorado, USA), belongs to the Boulder BO sublineage and was given to David Hirsh by Nigon’s student Jean-Louis Brun in 1983, and used in many of the original studies of TEs in *C. elegans* (Liao *et al*. 1983; Rosenzweig *et al*. 1983a; Moerman and Waterston 1984; Mori *et al*. 1988; Nigon and Felix 2017). A second Bergerac strain, RW6999, is relatively understudied and reported to be a subclone of RW7000 as per the Caenorhabditis Genetics Center (CGC) (https://cgc.umn.edu/strain/RW6999). A third strain, CB4851, belonging to the Cambridge sublineage (Nigon and Felix 2017), was shared with Sydney Brenner in 1969 (Hodgkin and Doniach 1997; Nigon and Félix 2017).

### Fitness assays for four life-history traits and statistical analyses of fitness data

To quantify the fitness of Bergerac strains RW7000, RW6999, and CB4851 relative to the laboratory strain Bristol N2 (also referred to as N2), we assayed four fitness-related, life-history traits, namely (i) productivity, (ii) survivorship to adulthood (also referred to as survivorship), (iii) longevity, and (iv) developmental rate as previously described (Katju *et al*. 2015, 2018; Dubie *et al*. 2020). All assays were conducted on Nematode Growth Medium (NGM) agar plates seeded with the *E. coli* strain OP50 at 20°C, the standard temperature for *C. elegans* culturing. Cryopreserved stocks of RW7000, RW6999, and CB4851 and N2 were thawed and individual worms isolated onto NGM plates. In order to establish independent lines, 15 and 20 worms were isolated for the wild type N2 and each Bergerac strain, respectively. After the establishment of multiple lines per strain, the worms were allowed to self and five of their L4 larval stage progeny were isolated singly onto a new plate, thereby establishing five sublines per line (total 15 × 5 = 75 for N2; 20 × 5 = 100 for each Bergerac strain). To negate the possibility of maternal or grandmaternal effects (Lynch 1985), each replicate was propagated by single-worm transfer for one more generation. This hierarchical structure (strains, lines, and sublines) combined with the fact that hermaphrodites self-fertilize to produce the next generation, minimizes genetic divergence among replicates, and allows for a clean estimate of environmental variance by comparison of sublines.

Three fitness assays (development, productivity, and longevity) were conducted on a single, third-generation worm isolated from each subline as an L1 larva. To measure the developmental rate, commencing 36h after the initial isolation of the L1 larva, each worm was checked every 2h to identify the time (in hours) until the first egg was visible in the worm’s uterus. Upon noting the presence of the first egg in the uterus, the worm was scored as having developed to adulthood. This initial measurement of the time (hours) from L1 larva to an egg-bearing adult yielded the developmental time. The inverse of the developmental time yielded the developmental rate. Worms that died before reaching adulthood were not scored.

To assay productivity, each worm that developed to adulthood was transferred to a new plate every 24h for eight consecutive days. After transferring the worm to a new plate, the eggs from the previous plate were allowed to hatch for an additional 24h, then stored at 4°C for a minimum of one month to allow the progeny to die without producing offspring. Counts were conducted by staining each plate with a 0.075% water dilution of toluidine blue, which temporarily makes the progeny stand out white against a purple background to facilitate visualization for worm counts.

Following eight days of daily transfers to a fresh plate as part of the productivity assay, each worm was transferred to a fresh NGM agar plate seeded with the *E. coli* OP50 until death in order to score longevity in days. In order to score longevity, the worms were monitored each day for movement and pharyngeal pumping. When no movement was detected, the agar pad near the worm was gently tapped. If no response was detected, the tail of the worm was tapped. If the worm failed to respond, it was recorded as dead and the days to mortality were noted.

To assay survivorship to adulthood, we used 10 L1 siblings of each third-generation worm assayed for the preceding three life-history traits. The 10 L1 larvae were isolated onto a seeded 60mm NGM agar plate. For some sublines, less than 10 L1 individuals were isolated due to the low and delayed productivity of Bergerac worms. 36h after isolation, the plates were checked for worms that survived to adulthood, and each plate was scored using the fraction of worms that survived to adulthood (values ranged from 0 to 1). For plates with desiccated worms on the edge of the plate, worms were still scored as surviving if eggs were observed in the uterus, to provide a more conservative estimate of survivorship.

For each of the four fitness traits, a two-level nested Model I ANOVA with unequal sample sizes (Sokal and Rohlf 1995) was employed to partition the total phenotypic variance into among- and within line-components. The highest level of classification tested for a treatment effect (four strains as treatments: (i) wild type control N2, (ii) strain RW7000, (iii) strain RW6999, and (iv) strain CB4851. The next level of hierarchy tested for a subgroup variance component (difference among independent lines within a treatment). The last hierarchical level estimated the subline variance. To conduct pairwise comparisons for all strain pairs per fitness trait, a Tukey-Kramer HSD test for unequal sample sizes was used at a 5% experiment-wise error rate.

### Motility Assays

In addition to the fitness assays, motility assays were conducted to quantify several locomotory traits in the Bergerac strains relative to the laboratory strain N2. Cryopreserved stocks of each strain were thawed and bottlenecked for three generations to establish five replicates for each strain, using procedures similar to the previously described fitness assays. Two identical assays were conducted on separate days, resulting in recordings of 10 plates for each strain in total. When the progeny of each third-generation replicate reached the L4 life stage, 10 hermaphrodite siblings were isolated onto a single 35×100 mm plate with 3.25 mL NGM. 21-22h after isolating L4s for each strain, the young adult worms were removed from a 20°C incubator and transferred to an unseeded NGM plate. Within 3 min of the transfer, a 60s video was recorded using Zen on an Axio Zoom.V16 Zoom microscope equipped with an AxioCam 503 Color camera at 5fps. The objective lens used for this analysis was the Zeiss PlanNeoFluar Z 1x/0.25 FWD 56mm. The conversion factor for videos provided by the Zen program was 25.94 *µ*M/pixel. The resolution of the final videos was 968×730. The order we recorded strains was randomized, and worms were kept in the incubator as long as possible to limit environmental variance. The temperature in the recording room was recorded at least three times for each assay and remained ∼21 ± 1°C. These details are provided to increase the reproducibility of our motility assay in a manner consistent with the suggestions of Angstman *et al*. (2016).

A custom Matlab program called Zentracker was used to analyze assay videos (https://github.com/wormtracker/zentracker). After inputting the scaling factor provided by the scope, all videos were tracked at intensities ranging from 130-180, then manually validated to ensure all worms were tracked and non-worm objects were removed. After checking the validity, the values for average speed, length, area and direction change were extracted for each video. In this analysis, averages were defined as the overall average value of all the valid datapoints.

### DNA extraction and Illumina sequencing

Genomic DNA was extracted from Bergerac and N2 control lines as previously described (Konrad *et al*. 2018) with libraries prepared using the Nextera DNAflex library kit (Illumina, San Diego, CA). Libraries were sequenced on the Illumina Novaseq6000 platform (2×150 bp) at the North Texas Genome Center at the University of Texas at Arlington.

### Tc1 copy-number estimation with ddPCR

Digital droplet polymerase chain reaction (ddPCR) copy-number variation (CNV) assay was performed following the Bio-Rad ddPCR Copy-Number Variation Assay protocol (https://www.bio-rad.com/webroot/web/pdf/lsr/literature/10033173.pdf) for lines N2, CB4851, RW6999, and RW7000. The ddPCR utilizing a Tc*1* targeting probe and a *daf-3* (a single copy reference) targeting probe with Fluorescein (FAM) were analyzed on the Bio-Rad QX ONE ddPCR system. The haploid genome-wide copy-number of Tc*1* was determined based on the estimated copy-number of Tc*1*-positive relative to the *daf-3* single-copy control. The reactions were run using two concentrations of template genomic DNA: (i) the first, at 0.1 ng/*µ*L, to allow a sufficient number of *daf-3* positive droplets, and (ii) another, a 100-fold dilution of 0.001 ng/*µ*L, to avoid saturation of positive Tc*1* droplets. The number of *daf-3* positives was then divided by the dilution factor to identify the proportion of negative droplets for Poisson calculations shown below. The cutoffs for positive and negative droplets were assigned manually on the QX Manager Software (1.2 Standard Edition) provided by Bio-Rad based on the separation of the droplets.

### Tc1 copy-number estimation from whole-genome sequence (WGS) data

Tc*1* copy-number in each genome was also estimated using the McClintock meta-pipeline, v2.0.0 (Nelson *et al*. 2017) which combines many TE-detection algorithms to identify reference and nonreference TE insertions in each genome. A standard consensus sequence of a 1,610 bp Tc*1* element was used as an input. Each TE-detection algorithm produces unique results containing both correct calls and false positives, with varying degrees of sensitivity and precision in simulated and real datasets (Nelson *et al*. 2017; Vendrell-Mir *et al*. 2019). McClintock v2.0.0 provides an estimate of normalized mean coverage. Five callers were chosen from the McClintock pipeline to estimate Tc*1* copy-number based on the absolute difference from the normalized mean coverage and ddPCR copy-number estimate: ngs_te_mapper2 (Han *et al*. 2021), RelocaTE (Robb *et al*. 2013), TEMP2 (Yu *et al*. 2021), RetroSeq (Keane *et al*. 2013), and TEFLoN (Adrion *et al*. 2017).

To visualize the overlap between the results of the various McClintock callers, the reference genome (PRJNA13758.WS279) was split into 1,000 bp windows using

BEDTools (Quinlan and Hall 2010). The BEDTools “intersect” command was used to create a file with a window for each non-reference Tc*1* call from each McClintock caller. Given that the reference Tc*1* elements tend to span several 1,000 bp windows, a similar procedure was used to assign reference calls to 100,000 bp windows, and subsequently these files were combined with the nonreference windows. Using a web Venn diagram tool (http://bioinformatics.psb.ugent.be/webtools/Venn/), Venn diagrams describing the overlap of Tc*1* calls for each method in each strain were produced, along with a Venn diagram describing the overlap of TEMP2 Tc*1* calls between the Bergerac strains.

In addition to Tc*1*, we employed the TEMP2 caller from McClintock to estimate copy-number for other TEs, namely *Tc2*, *Tc3*, *Tc4v*, *Tc5*, *Tc6*, *Tc9*, *TURMOIL1*, *TURMOIL2*, *MARINER2*, *MARINER3*, *MARINER4*, and *MARINER5*.

### Tc1 landing site sequence analysis

Due to the high positional accuracy of the TEMP2 caller (∼0.90) in the prediction of synthetic TE insertions used as benchmarks in the original McClintock publication, it was chosen for analyses of TE landing sites (Nelson *et al*. 2017). The sequence context of each Tc*1* insertion predicted by TEMP2 was analyzed in each Bergerac strain. The FASTQ reads generated by Illumina whole-genome sequencing were aligned to the *C. elegans* reference genome (PRJNA13758.WS279) with the Burrows-Wheeler Aligner (BWA), version 0.7.12-r1039. Using the BED file generated by the McClintock pipeline, sequences ± 25 bp from each insertion site were extracted from the BWA alignments.

These sequences were aligned using ClustalW and a consensus sequence was generated for each strain with IUPAC nucleotide code by analyzing the distribution of nucleotides ± 6 bp from the center of the insertion.

### Statistical analyses of TE distribution in the Bergerac genomes

To assess the environment of Tc*1* landing sites within the Bergerac strains’ genomes, genomic coordinates specific to various genomic features were used to define the genomic context of Tc*1* insertions. Annotation files were created for chromosomal arms, cores, and tips, exon, intron, or intergenic regions, as well as various histone modification environments. To preclude overlapping regions with the annotation files from yielding multiple counts, these annotation files were then merged to prevent overlapping regions within the annotation files from yielding multiple counts using the “merge" command of the BEDTools software package (Quinlan and Hall 2010). Finally, the merged BED files denoting the landing site environment and the resulting Tc*1* BED files for each line were analyzed for overlap using the bedtools “intersect" command. For exon and intron overlap with identified Tc*1* sites, the merged exon and intron annotations were analyzed for intersections with the “-v” flag ensuring only intron exclusive genomic sites used to call intronic sites. The resulting counts were used as input for chi-squared tests, using the proportion of genomic coverage to calculate expected values within the R statistical analysis software (R Core Team 2014).

### Genes with exons disrupted by Tc1

Using the *C. elegans* N2 reference annotations file from WormBase (version WS279; www.wormbase.org), a table of genes with predicted exonic or intronic Tc*1* insertions according to TEMP2 was generated along with a list of intergenic insertions. Some insertions in this table are repeated, as the exons of genes with multiple transcripts do not always agree. Using the WormBase tool SimpleMine (https://wormbase.org/tools/mine/simplemine.cgi), a table containing a description of RNAi phenotypes, allele phenotypes, concise descriptions, and automated descriptions for genes disrupted by Tc*1* was created.

### Analysis of SNPs and CNV in the Bergerac strains

In order to identify candidate mutations responsible for Tc*1* proliferation in the Bergerac strains, the MiModd package (www.celegans.de/mimodd) was used to identify SNPs, indels, and deletions in the three Bergerac strains, the N2 strain housed in the laboratory, and seven additional natural isolates: AB1, ECA243, JU1568, JU2565, JU394, NIC1049, and NIC2. One isolate, ECA243, is a renamed sample of the strain CB4851 that was previously sequenced. The BAM alignments for the seven chosen natural isolates were obtained from the CeNDR database, release 20210121 (Cook *et al*. 2017). The reads for the Bergerac strains and our laboratory sample of N2 were cleaned with Fastp using default settings and aligned with BWA-MEM (Burrows-Wheeler Aligner maximal exact match) to the *C. elegans* reference genome PRJNA13758.WS276 to replicate the CeNDR procedure. All 11 BAM files were then merged, and MiModd was used to extract SNPs and indels for all strains. These variants were filtered to keep homozygous variants with a minimum depth of 3. SNPeff, a tool to predict the effect of SNPs on protein-coding genes, was used to annotate these variants (Cingolani *et al*. 2012). In addition to SNPeff, SIFT 4G was used to predict which amino acid changes in the Bergerac strains are likely to be deleterious to protein function. Deletions were also called for all strains using MiModd, with a maximum coverage of 4 and a minimum size of 100 bp. After calling variants and deletions, the mutations were filtered to retain homozygous variants identified in all three Bergerac strains and absent in all non-Bergerac strains. Genes with deletions were determined using the *C. elegans* WS276 reference gff3 file. Summary tables for all mutated genes were created using SimpleMine. Finally, to place the Bergerac strains in a phylogenetic context, we ran a maximum likelihood analysis with PhyML (Guindon *et al*. 2010), using a GTR substitution model with a discrete gamma distribution and four rate categories on 11 *C. elegans* strains, which included the three Bergerac strains and N2.

## RESULTS

### Significant variation in Tc1 copy-number among the Bergerac strains

The standard laboratory strain of *C. elegans*, Bristol N2, has diverged genetically since it was first isolated through the independent accumulation of base substitutions, copy-number changes and TE activity during propagation in different laboratories (Lipinski *et al*. 2011). The estimated number of Tc*1* elements in low Tc*1* copy-number strains, such as Bristol N2 typically ranges from 20-30 copies (Emmons *et al*. 1983; Liao *et al*. 1983; Egilmez *et al*. 1995). The median and average number of Tc*1* elements in our reference Bristol N2 was 28, with a range of 26-30 copies based on five methods (ngs_te_mapper 2, RelocaTE, TEMP2, Teflon, Normalized TE coverage) implemented in the McClintock pipeline (**Figure 1A, Table 1, Supplemental Table S1**). The RetroSeq caller of the McClintock program does not call non-reference TEs; hence an estimate was omitted for the N2 strain. ddPCR results from our N2 control estimated the Tc*1* copy-number as 29.

**Figure 1.**
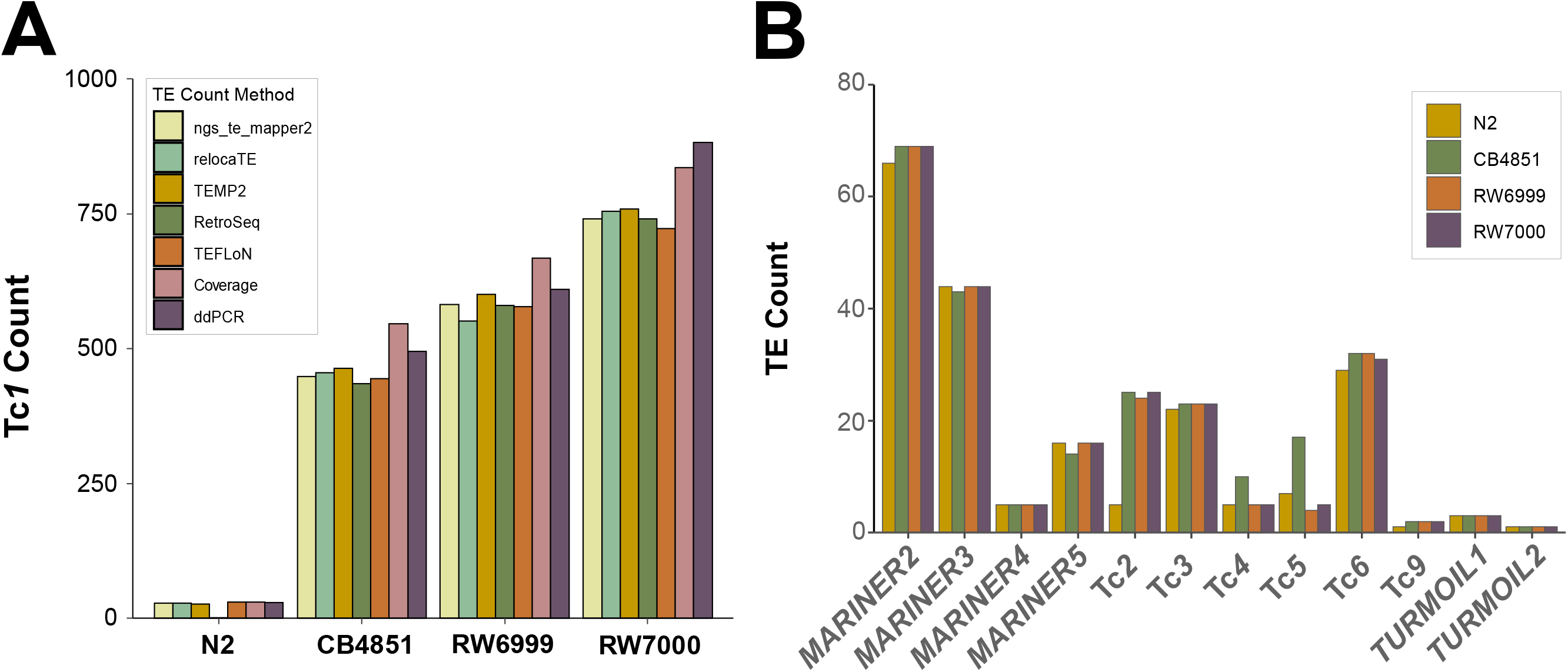
Proliferation of the Tc*1* transposon in the Bergerac strains of *C. elegans*. (A) Counts of the Tc*1* transposon in our laboratory isolate of Bristol N2 and three Bergerac strains, CB4851, RW6999, and RW7000 using various computational and molecular methods. Computational methods included were ngs_te_mapper, RelocaTE, TEMP2, RetroSeq, TEFLoN, and coverage. Copy-number estimation via ddPCR was the sole molecular method. (B) Counts of other/non-Tc*1* transposable elements are displayed for N2, CB4851, RW6999, and RW7000. Tc*2* was the only transposable element with a marked increase in the three Bergerac lines compared to N2, while Tc*4* and Tc*5* were found to be marginally higher in CB4851.

**Table 1.**
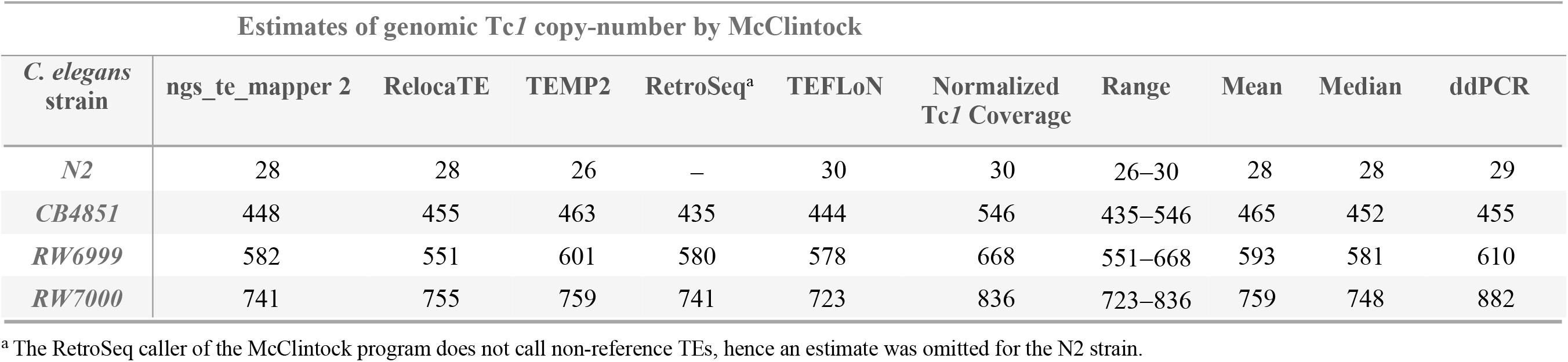
Estimated number of Tc1 transposable elements in the reference N2 and three Bergerac strains of C. elegans. . The haploid number of Tc*1* as determined by several TE element callers in the McClintock pipeline (Nelson *et al*. 2017). The range, means and medians are calculated from estimates provided by six selected callers in the McClintock pipeline. Copy-number estimates of Tc*1*from ddPCR are included for comparison.

The three Bergerac strains display substantial variation in Tc*1* copy-number that is consistent between different variant callers notwithstanding some variation in results between different methods (**Figure 1A, Table 1**). The range in Tc*1* copy-number estimates among the Bergerac strains was non-overlapping (**Table 1**). We first estimated Tc*1* copy-number in the three Bergerac strains by employing six computational methods/callers available in the McClintock pipeline (ngs_te_mapper 2, RelocaTE, RetroSeq, TEMP2, TEFLoN, and Normalized TE coverage). CB4851 harbored the lowest number of Tc*1* elements with a mean and median of 465 and 451, respectively. The mean and median of Tc*1* copy-number in RW6999 was 593 and 581, respectively. RW7000 has the highest abundance of Tc*1* elements with a mean and median estimate of 759 and 748, respectively (**Figure 1B, Table 1**). The estimates based on normalized Tc*1* coverage were 12-21% greater than those based on the five methods that use paired-ends and split reads to estimate copy-number. BED files containing the locations of all calls produced by McClintock are provided in **Supplemental Table S1**. ddPCR estimates of Tc*1* copy-number in the three Bergerac strains were consistently higher than their corresponding median copy-number estimates from McClintock, ranging from 495, 610, and 882 copies for CB4851, RW6999 and RW7000, respectively. Both ddPCR and McClintock estimates support the general conclusion that the three Bergerac strains can vary considerably with respect to Tc*1* copy-number.

To assess whether McClintock component methods were calling Tc*1* insertions in similar locations, Tc*1* calls produced by ngs_te_mapper2, RelocaTE, TEMP2, RetroSeq, and TEFLoN were assigned to 1,000 bp windows and compared (**Figure 2**). The vast majority of Tc*1* calls are supported by multiple callers, with 70% of the 523 unique Tc*1* calls in CB4851 supported by all five callers, and 83% of calls supported by four or more callers. In addition to the comparison of Tc*1* insert locations between callers, the overlap of insert locations between the Bergerac strains was assessed by comparing the locations of calls produced by TEMP2 (**Supplemental Figure S1**). Though a fraction of insert sites are shared between all three strains, the Bergerac strains display high variability in the location of insert sites, with large differences observed between the strain CB4851 and the two RW strains. This pattern reflects continued Tc*1* mobility and proliferation during the separate laboratory cultivations of these strains reported by Nigon and Felix (2017), with the RW strains sharing a more recent common ancestor.

**Figure 2.**
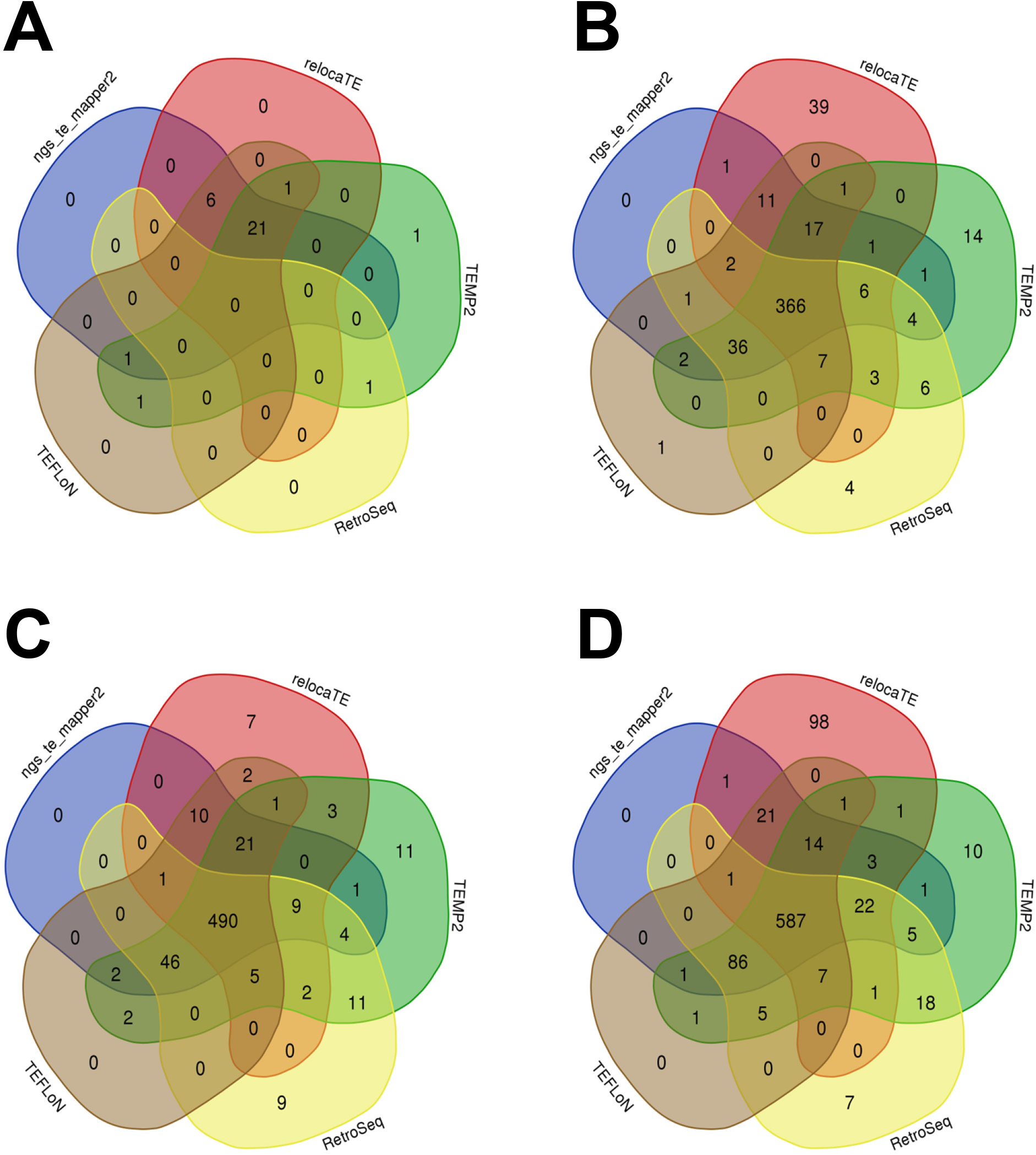
Comparison of Tc*1* insertions predicted by McClintock component methods. The positional agreement of the McClintock component methods was assessed by assigning Tc*1* calls to windows (1,000 bp and 100,000 bp windows for non-reference and reference calls, respectively) followed by a comparison of the calls made by each component method. The Venn diagrams compare the agreement of Tc*1* calls for (A) N2, (B) CB4851, (C) RW6999, and (D) RW7000. Computational methods included were ngs_te_mapper2 (blue), RelocaTE (red), TEMP2 (green), RetroSeq (yellow), and TEFLoN (brown).The vast majority of Tc*1* insertions were called by multiple methods.

### Variation in non-Tc1 elements in the Bergerac strains

The McClintock package was also used to estimate the copy-number of a wider variety of TEs in the three Bergerac strain relative to N2 (**Supplemental Table S1**). There were no differences in the estimated counts between N2 and the Bergerac strains for the majority of non-Tc*1* elements (*Mariner2-5,* Tc*3,* Tc*6,* Tc*9, Turmoil1* and *2*) (**Figure 1B**). However, the Bergerac strains appear to have more Tc*2* elements than N2. The average estimated count of Tc*2* in was three and 20 in N2 and the Bergerac strains, respectively. Coverage estimates using normalized read depth ranged between 25−30 in the Bergerac strains and four in N2. The average counts of Tc*4* and Tc*5* appeared to be higher in CB4851 than in the other two Bergerac strains (RW6999 and RW7000) and N2. However, the results were highly variable between different callers in McClintock, especially for Tc*5* where the number of counts ranged from 0−40, and the distribution of counts exhibited considerable overlap between the four strains. Furthermore, normalized read depth did not provide support for differences in Tc*5* counts between the strains, with estimates of 13−14 elements per strain.

### Bergerac strains have significantly lower fitness than the wild type N2 strain

Given that the Bergerac strain was one of the earliest *C. elegans* isolates to be cultured under laboratory conditions (Nigon and Felix 2017), preceding studies have described its temperature-sensitivity (Fatt and Dogherty 1963), low fertility (Abdulkader and Brun 1980; Lee *et al*. 2016), and other phenotypic abnormalities (Hodgkin and Doniach 1997). This study aimed to empirically quantify the fitness of the three Bergerac strains (CB4851, RW6999, RW7000) relative to the Bristol N2 strain by measuring four fitness traits, namely productivity, survivorship to adulthood, longevity, and developmental time. The phenotypic fitness assays comprised measurements on five replicates each, where possible, of 15 N2 lines and 20 lines each of the three Bergerac strains. The mean fitness values for each measured trait are displayed in **Figures 3A-D** and **Table 2**.

**Figure 3.**
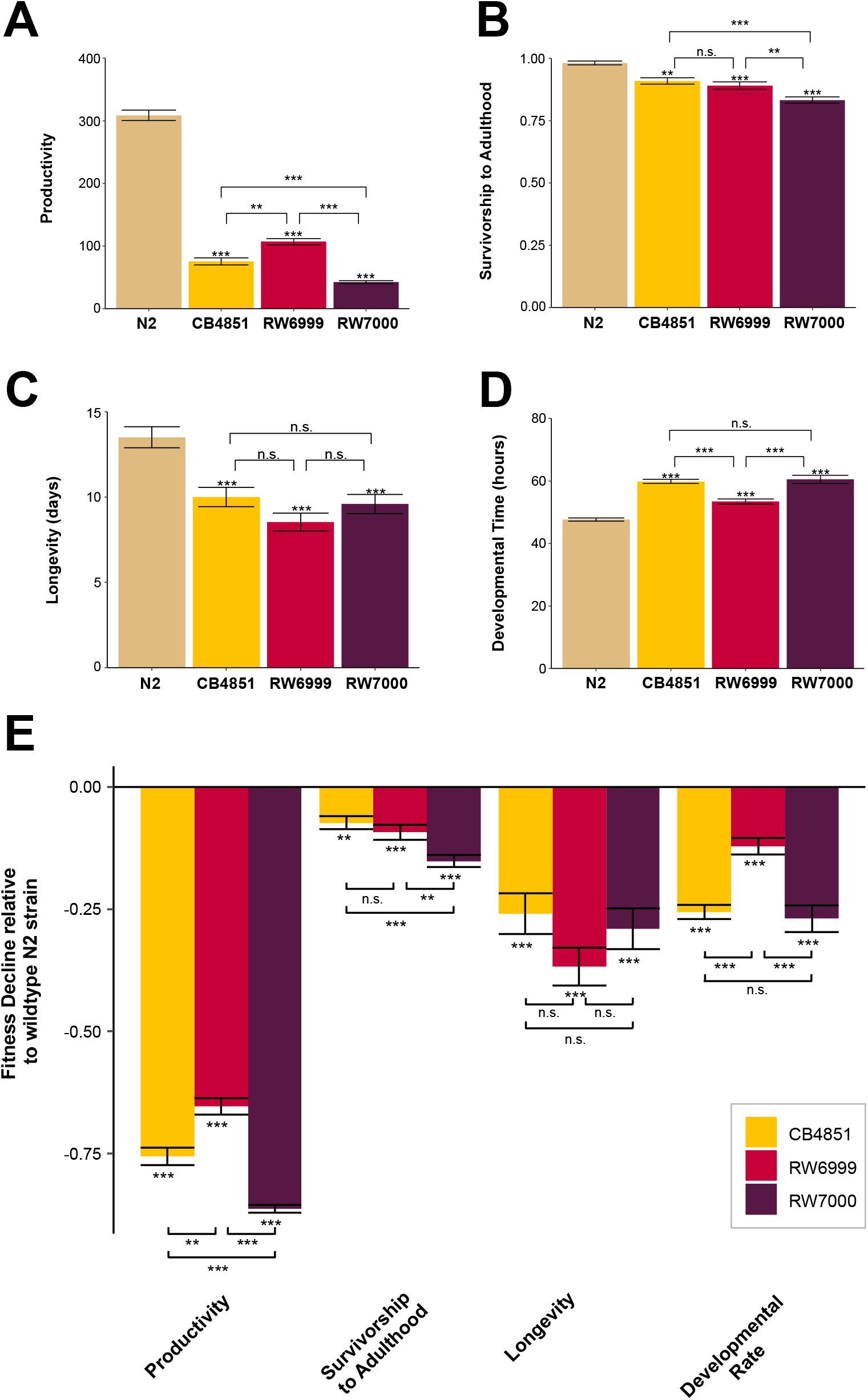
Significant fitness reduction in Bergerac strains relative to N2. Means for four fitness-related traits, namely (A) productivity, (B) survivorship to adulthood, (C) longevity, and (D) developmental time observed in the N2 control, and the three Bergerac strains CB4851, RW6999, and RW7000. (E) Decline in mean relative fitness of the three Bergerac strains relative to the N2 strain. For simplicity, the mean relative fitness value for each of the four traits in the Bristol N2 control was scaled to a value of 1 (not displayed). For all panels, error bars represent one standard error. Significance was determined via ANOVA and is displayed as asterisks where *p* ≤ 0.05*/0.01**/0.001***. Exact *p*-values for the pairwise strain comparisons using Tukey-Kramer HSD can be found in Supplemental Tables S2-S5.

**Table 2.** Fitness of three C. elegans Bergerac strain relative to control worms of the laboratory strain, N2. For the N2 control, 15 lines each with five replicates were established (maximum *n* = 75). For each of the three Bergerac strains (CB4851, RW6999, and RW7000), we assayed 20 lines with five replicates each (maximum *n* = 100). Estimates of the mean phenotype for four fitness-related traits are provided for each of the four strains.

| *C. elegans* strain | Productivity | Survivorship to Adulthood (%) | Longevity (days) | Developmental Time (hours) |
| --- | --- | --- | --- | --- |
| N2 CONTROL | 309 | 98 | 14 | 48 |
| CB4851 | 75 | 91 | 10 | 60 |
| RW6999 | 107 | 89 | 9 | 53 |
| RW7000 | 43 | 83 | 10 | 60 |

**Table 3.** Two-level nested ANOVA for productivity, survivorship to adulthood, longevity, and developmental time of N2 control and Bergerac strains CB4851, RW6999, and RW7000.

| Source of variation | df | SS | MS | $F_s$ | $F_s'$ |
| --- | --- | --- | --- | --- | --- |
| ***Productivity*** |  |  |  |  |  |
| Among groups | 3 | 3069207 | 1023069 | 403.83 | 309.93**** |
| Among lines | 69 | 217546 | 3153 | 1.25 |  |
| Within lines (error) | <u>224</u> | 567480 | 2533 |  |  |
| Total | 296 |  |  |  |  |
| ***Survivorship to Adulthood*** |  |  |  |  |  |
| Among groups | 3 | 0.945 | 0.315 | 23.88 | 16.80**** |
| Among lines | 69 | 1.289 | 0.019 | 1.42* |  |
| Within lines (error) | <u>274</u> | 3.615 | 0.019 |  |  |
| Total | 346 |  |  |  |  |
| ***Longevity*** |  |  |  |  |  |
| Among groups | 3 | 1044 | 348.0 | 12.77 | 12.49**** |
| Among lines | 68 | 2029 | 29.8 | 1.10 |  |
| Within lines (error) | <u>220</u> | 5995 | 27.3 |  |  |
| Total | 291 |  |  |  |  |
| ***Developmental Time*** |  |  |  |  |  |
| Among groups | 3 | 7804 | 2601.3 | 35.51 | 34.06**** |
| Among lines | 69 | 5348 | 77.5 | 1.06 |  |
| Within lines (error) | <u>223</u> | 16337 | 73.3 |  |  |
| Total | 295 |  |  |  |  |

\* significance level of 0.05
\*\* significance level of 0.01
\*\*\* significance level of 0.001
\*\*\*\* significance level of 0.0001

The mean productivity of the N2, CB4851, RW6999, and RW7000 strains was 309, 75, 107, and 43 offspring, respectively (**Figure 3A; Table 2**). The three Bergerac strains exhibited a 65-86% reduction in mean productivity relative to the N2 strain (**Figure 3E**). As reported in a previous study of Bergerac-derived recombinant inbred lines (RILs) (Shook and Johnson 1999), many Bergerac worms were observed to die by matricidal hatching (“worm bagging” or “bagging”), a phenotype characterized by larvae hatching within the hermaphrodite before egg-laying occurs. However, these worms were observed to lay some eggs before bagging and were included in the analysis to provide a realistic estimate of mean productivity in laboratory conditions. ANOVA analyses found a significant variance component for productivity among the four strains (*F_S_* ^′^ = 309.93; *p* < 0.0001) whereas among-line divergence was nonsignificant (**Table 3**).

The mean survivorship to adulthood of the N2, CB4851, RW6999, and RW7000 strains was 98, 91, 89, and 83%, respectively (**Figure 3B; Table 2**). The three Bergerac strains exhibited a 7-15% reduction in mean survivorship relative to the N2 strain (**Figure 3E**). ANOVA analyses found a significant variance component for survivorship among the four strains (*F_s_*^′^ = 16.8; *p* < 0.0001) as well as among-line divergence (*F_s_* = 1.42; *p* < 0.05) (**Table 3**). As was observed for productivity, RW7000 exhibited the lowest survivorship among the three Bergerac strains relative to N2.

The mean longevity of the N2, CB4851, RW6999, and RW7000 strains was 14, 10, 9, and 10 days, respectively (**Figure 3C; Table 2**). The three Bergerac strains exhibited a 29-36% reduction in longevity relative to the N2 strain (**Figure 3E**). ANOVA analyses found a significant variance component for longevity among the four strains (*F_s_* = 12.49; *p* < 0.0001) whereas among-line divergence was nonsignificant (**Table 3**).

The mean developmental time of the N2, CB4851, RW6999, and RW7000 strains was 48, 60, 53, and 60 hours, respectively (**Figure 3D; Table 2**). Hence, all three Bergerac strains exhibited delayed development to reproductive maturity relative to N2 (10-25% longer). ANOVA analyses found a significant variance component for developmental time among the four strains (*F_s_* ^′^ = 34.06; *p* < 0.0001) whereas among-line divergence was nonsignificant (**Table 3**). We further calculated the developmental rate as the inverse of the relative developmental time. Relative to N2, the three Bergerac strains exhibited a 9-20% reduction in their developmental rate (**Figure 3E**), with RW6999 and RW7000 exhibiting the greatest reduction in developmental rate.

The fitness assays additionally detected significant differences among Bergerac strains for all fitness-related traits. We conducted comparisons among pairs of means based on unequal sample sizes using the Tukey–Kramer HSD method to determine which strain pairs differed significantly from each other for the four assay fitness traits (**Figures 3A-E, Supplemental Tables S2-S5**). All three Bergerac strains were significantly different from N2 for each of the four fitness traits. With respect to productivity, all pair-wise strain comparisons were significant with the relationship expressed as follows: RW7000 > CB4851 > RW6999 > N2 (**Table 2, Supplemental Table S2**). With the exception of the CB4851 *vs.* RW6999 comparison, all other pair-wise strains comparisons showed significant differences for survivorship to adulthood (RW7000 < RW6999, *p* = 4.70 × 10^-3^; RW7000 < CB4851, *p* = 1.90 × 10^-4^) N2 (**Table 2, Supplemental Table S3**). Survivorship to adulthood of the four strains showed the following trend: RW7000 > RW6999/CB4851 > N2. While all of the three Bergerac strains had significantly reduced mean longevity relative to N2, they did not differ significantly from each other with all Bergerac strains surviving an average of 9-10 days after the L1 larval stage N2 (**Table 2, Supplemental Table S4**). With the exception of the CB4851 *vs.* RW7000 comparison, all other pair-wise strains comparisons showed significant differences for mean developmental time (RW6999 < RW7000, *p* = 1.87 × 10^-6^; RW6999 < CB4851, *p* = 5.94 × 10^-5^). (**Table 2, Supplemental Table S5**). Developmental time of the four strains showed the following trend: N2 > RW6999 > RW7000 > CB4851/RW7000.

### Phenotypic divergence of Bergerac strains with respect to motility- and size-related traits

The motility assays revealed variable levels of phenotypic divergence for several locomotory and size-related traits in the Bergerac strains. We first analyzed the four traits (speed, direction, body length and body area) in the wild type N2 strain. Our analysis of the average speed and direction change of N2 worms was concordant with the values observed by Angstman *et al*. (2006), despite our use of a different worm tracking program. Our N2 average speed was 177.8 *μ*m/*s* with an SD of 48.9 *μ*m/*s* (SE 15.5) compared to an average 146.9 *μ*m/*s* with an SD of 55.3 *μ*m/*s* measured by Angstman *et al*. (2006). Similarly, our N2 average direction change was 0.531 radians/*s* with an SD of 0.218 radians/*s* (SE 0.069) compared to Angstman *et al*.’s (2006) average direction change of 0.70 radians/*s* with an SD of 0.30 radians/*s*, after converting their measurements from degrees to radians. Similarly, the average length of our N2 worms (799 *μ*m, SD 56.2 *μ*m, SE 17.8) is close to a previously reported length of N2 after the fourth larval molt, 850 *μ*m (Byerly *et al*. 1976). Trait means for speed, body length, body area, and direction change are reported in **Figures 4A-D** and **Table 4**. As each data point represents the average of ten worms on a plate for 1min, a one-level ANOVA test was used to reveal significant differences for each trait measured (speed, direction, body length, and body area). We additionally conducted comparisons among pairs of means using the Tukey–Kramer HSD method to determine which strain pairs differed significantly from each other for the four motility- and size-associated traits (**Figures 4A-E, Supplemental Tables S6-S9**).

**Figure 4.**
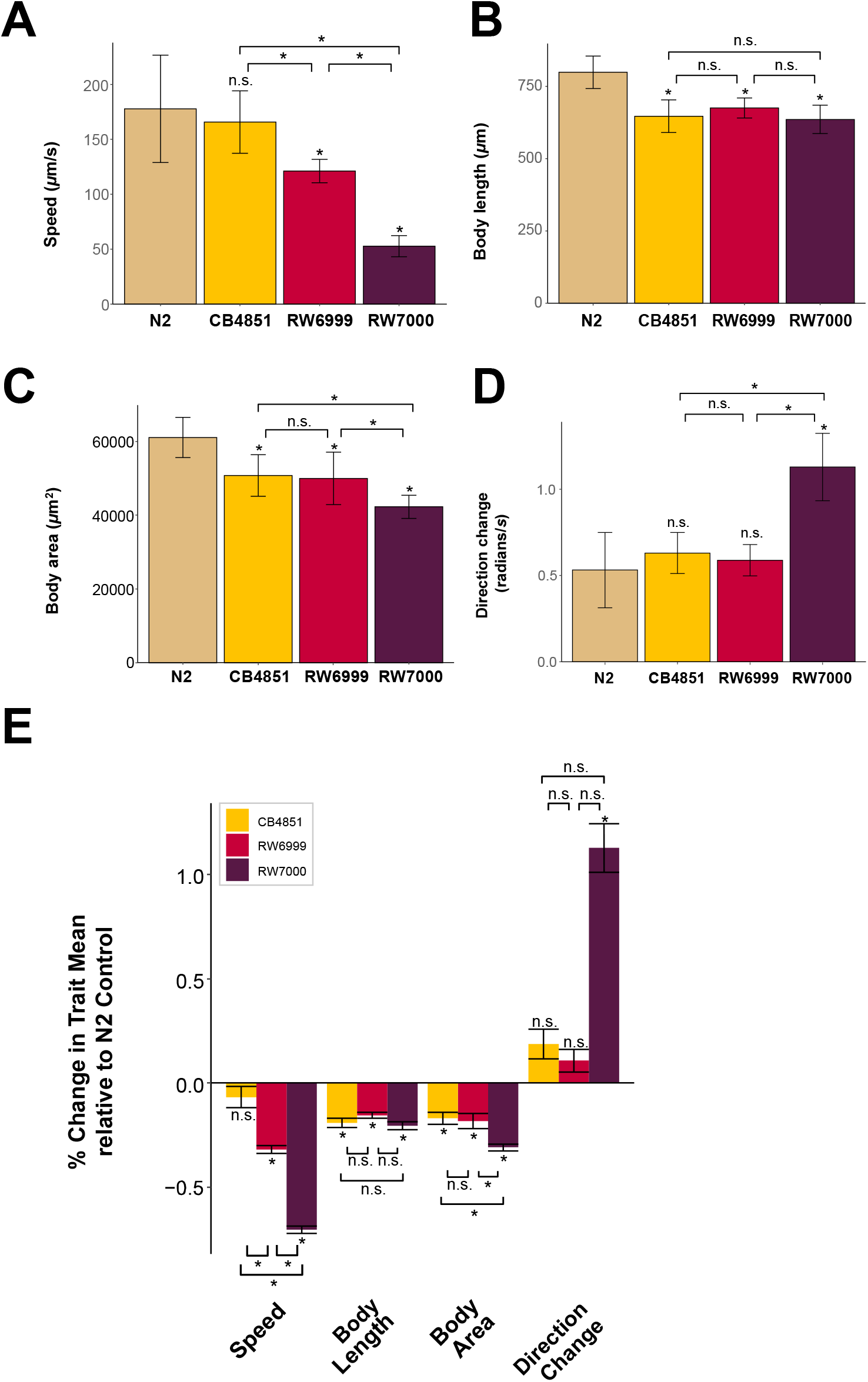
Physical size reduction and aberrant behavior of the Bergerac strains. Mean values for four traits measured in the motility analyses, namely (A) speed, (B) body length, (C) body area, (C), and direction change (D) in the N2 control, and the three Bergerac strains CB4851, RW6999, and RW7000. (E) Mean relative changes between the three Bergerac strains and the N2 control. For simplicity, the mean relative value for each of the four traits in the Bristol N2 control was scaled to a value of 1 (nor displayed). The stars on brackets summarize *p*-values for Tukey-Kramer HSD comparisons between Bergerac strains. Stars on top of bars reflect *p-*values for Tukey-Kramer HSD comparisons to N2. Exact *p*-values for the pairwise strain comparisons using Tukey-Kramer HSD can be found in Supplemental Tables S6-S9.

**Table 4.** Motility- and size-associated traits of three C. elegans Bergerac strain relative to control worms of the laboratory strain, N2. Traits measures for each of the four strains are for *n* = 100 worms across 10 plates (10 worms/plate). Mean values for all four motility-related traits are provided for each of the four strains.

| *C. elegans*<br>strain | Speed<br>($\mu\text{m/s}$) | Length<br>($\mu\text{m}$) | Area<br>($\mu\text{m}^2$) | Direction Change<br>(radians/s) |
| --- | --- | --- | --- | --- |
| *N2 control* | 178 | 799 | 61,078 | 0.53 |
| *CB4851* | 166 | 647 | 50,758 | 0.63 |
| *RW6999* | 121 | 675 | 49,966 | 0.59 |
| *RW7000* | 53 | 636 | 42,266 | 1.13 |

The mean speed of the N2, CB4851, RW6999, and RW7000 strains was 178, 166, 121, and 53 *m*m/*s*, respectively (**Figure 4A; Table 4**). The three Bergerac strains exhibited ∼7-70% reduction in speed relative to the N2 strain (**Figure 4E**). ANOVA analyses found a significant variance component for speed among the four strains (*F_s_* = 37.57; *p* = 3.50 × 10^-11^) (**Table 5**). With the exception of the CB4851 *vs.* N2 comparison, all other pair-wise strains comparisons showed significant differences for mean speed (RW7000 < CB4851, *p* = 1.51 × 10^-9^; RW7000 < RW6999, *p* = 4.14 × 10^-5^; RW6999 < CB4851, *p* = 8.30 × 10^-3^) (**Table 4, Supplemental Table S6**). Strain speed showed the following trend: N2/CB4851 > RW6999 > RW7000.

**Table 5.** ANOVA analyses for four motility and size-associated traits in wild type N2 and three Bergerac strains (CB4851, RW6999, and RW7000) of *C. elegans*.

| Source of variation | *df* | SS | MS | *F<sub>s</sub>* |
| --- | --- | --- | --- | --- |
| ***Speed (μm/s)*** |  |  |  |  |
| Among groups | 3 | 96109 | 32036 | 37.57*** |
| Within lines (error) | <u>36</u> | <u>30697</u> | 852 |  |
| Total | 39 | 126807 |  |  |
| ***Length (μm)*** |  |  |  |  |
| Among groups | 3 | 169232 | 56410 | 22.58**** |
| Within lines (error) | <u>36</u> | <u>89950</u> | 2498 |  |
| Total | 39 | 259182 |  |  |
| ***Area (μm<sup>2</sup>)*** |  |  |  |  |
| Among groups | 3 | 1789912485 | 596637495 | 19.59**** |
| Within lines (error) | <u>36</u> | <u>1096184461</u> | 30449568 |  |
| Total | 39 | 2886096946 |  |  |
| ***Direction Change (radians/s)*** |  |  |  |  |
| Among groups | 3 | 2.288 | 0.763 | 28.15**** |
| Within lines (error) | <u>36</u> | <u>0.976</u> | 0.027 |  |
| Total | 39 | 3.264 |  |  |

\* significance level of 0.05
\*\* significance level of 0.01
\*\*\* significance level of 0.001
\*\*\*\* significance level of 0.0001

The mean body length of the N2, CB4851, RW6999, and RW7000 strains was 799, 647, 675, and 636 *m*m, respectively (**Figure 4B; Table 4).** The three Bergerac strains exhibited ∼16-20% reduction in body length relative to the N2 strain (**Figure 4E**). ANOVA analyses found a significant variance component for body length among the four strains (*F_s_* = 22.58; *p* = 2.14 × 10^-8^) (**Table 5**). While all of the three Bergerac strains had significantly reduced mean body length relative to N2, they did not differ significantly from each other (**Table 4, Supplemental Table S7**). Hence, body length showed the following trend: N2 > CB4851/RW6999/RW7000.

The mean body area of the N2, CB4851, RW6999, and RW7000 strains was 61078, 50758, 49966, and 42266 *m*m^2^, respectively (**Figure 4C; Table 4**). The three Bergerac strains exhibited ∼17-31% reduction in body area relative to the N2 strain (**Figure 4E**). ANOVA analyses found a significant variance component for body area among the four strains (*F_s_* = 19.59; *p* = 1.06 × 10^-7^) (**Table 5**). All three Bergerac strains had significantly reduced mean body area relative to N2 (**Supplemental Table S8**). The three Bergerac strains also exhibited some significant differences amongst each other with respect to body area. While the CB4851/RW6999 pair comparison was nonsignificant, RW7000 had significantly smaller body area relative to both CB4851 (*p* = 7.73 × 10^-3^) and RW6999 (*p* = 1.78 × 10^-2^) (**Supplemental Table S8**). Body area showed the following trend: N2 > CB4851/RW6999 > RW7000. Because all worms in these motility assays were approximately the same age, these size differences could be explained by the previously observed variation in developmental time. A Pearson test between the average developmental time for each strain and body area confirms that these variables are correlated (*p* = 1.17 × 10^-6^).

The mean direction change of the N2, CB4851, RW6999, and RW7000 strains was 0.53, 0.63, 0.59, and 1.13 radians/*s*, respectively (**Figure 4D; Table 4**). The three Bergerac strains exhibited ∼11-113% increase in direction change relative to the N2 strain (**Figure 4E**). ANOVA analyses found a significant variance component for direction change among the four strains (*F_s_* = 28.15; *p* = *p* = 1.50 × 10^-9^) (**Table 5**).

However, only RW7000 was significantly different from the three other strains (N2, and the other two Bergerac strains) with respect to direction change N2 (**Supplemental Table S9**). The average direction change of strain RW7000 was much larger than the other Bergerac strains (RW6999 < RW7000, *p* = 6.70 × 10^-8^; CB4851 < RW7000, *p* = 3.72 × 10^-7^). The reason for the large difference in the average direction change displayed by RW7000 is also visually apparent. While wild-type worm locomotion occurs in an approximately sinusoidal trajectory, the low fitness RW7000 worms tend to lay still in a much straighter orientation than other strains with shallower wave magnitude during their sinusoidal locomotion. Because their ability to move forward and turn seems to be restricted, any changes in movement tend to be in back-and-forth motion close to π radians, while other strains tend to move forward and make slow turns. Direction change showed the following trend: N2/CB4851/RW6999 < RW7000.

### Local sequence context of Tc1 insertion sites

Tc*1* had been well-documented to insert into 5′−TA−3′ target sites with a lightly conserved A/T rich motif (Rosenzweig *et al*. 1983b; Eide and Anderson 1988). Korswagen *et al*. (1996) analyzed 83 Tc*1* insertion sites to conclude a symmetric consensus sequence for Tc*1* insertion, namely CAYA**TA**TRTG. We further analyzed 1,659 Tc*1* insertion sites in the three Bergerac strains to derive a consensus insertion motif (**Figure 5**), confirming the conclusions of preceding studies (Eide and Anderson 1988; Korswagen *et al*. 1996). All the Tc*1* insertions were located at a 5′−TA−3′ target site within the consensus sequence. The consensus sequence was identified at 46,630 locations in the N2 reference genome (PRJNA13758.WS279).

**Figure 5.**
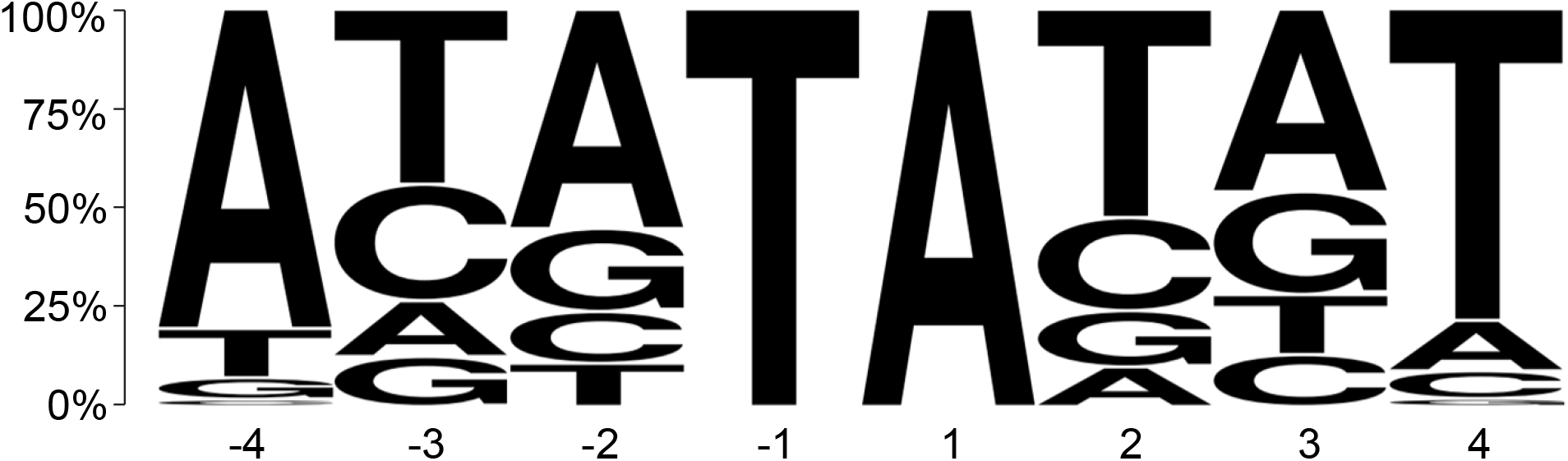
The Tc*1* insertion motif in the Bergerac strains. The vertical axis indicates the proportion of each base at four upstream and downstream positions from the insertion site. The motif for Tc*1* insertion sites was based on 1,659 insertions in this study and matched that presented in previous literature. Tc*1* invariably inserts between a T(-1) and an A(+1) with additional conserved A/T bases at positions -4 and +4.

### Nonrandom genomic distribution of Tc1 elements in the Bergerac strains

To determine whether Tc*1* proliferation in the Bergerac strains was random or influenced by genomic context, we classified the *C. elegans* genome based on four broad categories: (i) chromosomes, (ii) recombination domains, (iii) chromatin environment, and (iv) exonic, intronic or intergenic regions.

There is significant variation in chromosomal distribution of Tc*1* elements between the three Bergerac strains (*G* = 96.27, *p* = 3.33 × 10^-16^) (**Figures 6A and 6B**). This is primarily due to the relatively low number of Tc*1* elements on Chr. V in CB4851. When Chr. V was excluded from the analysis, the chromosomal distribution of Tc*1* was not significantly different between RW6999 and RW7000 (*G* = 1.07, *p* = 0.96), nor between the three Bergerac strains (*G* = 4.32, *p* = 0.83). Additionally, Tc*1* insertions were nonrandomly distributed across the six chromosomes (five autosomes and X) in the three Bergerac strains once corrected for chromosome length. Tc*1* elements were overrepresented on Chr. V and X in the closely-related strains RW6999 (*χ*^2^ = 49.06, *p* = 2.16 10^-9^) and RW7000 (*χ*^2^ = 50.88, *p* = 9.15 10^-10^). This Tc*1* overrepresentation was particularly pronounced on chromosome V which had 48% and 40% more Tc*1* elements than expected based on chromosome length alone in strains RW6999 and RW7000, respectively. When the number of 8 bp consensus Tc*1* insertion sites (5′−AYATATRT−3′) per chromosome is taken into account, the number of Tc*1* are 56% and 49% greater on chromosome V than expected in RW6999 and RW7000, respectively. Tc*1* number was better correlated with chromosome size (*R*^2^ = 0.96 for RW6999) than the number of consensus Tc*1* insertion sites per chromosome (*R*^2^ = 0.70 for RW6999).

**Figure 6.**
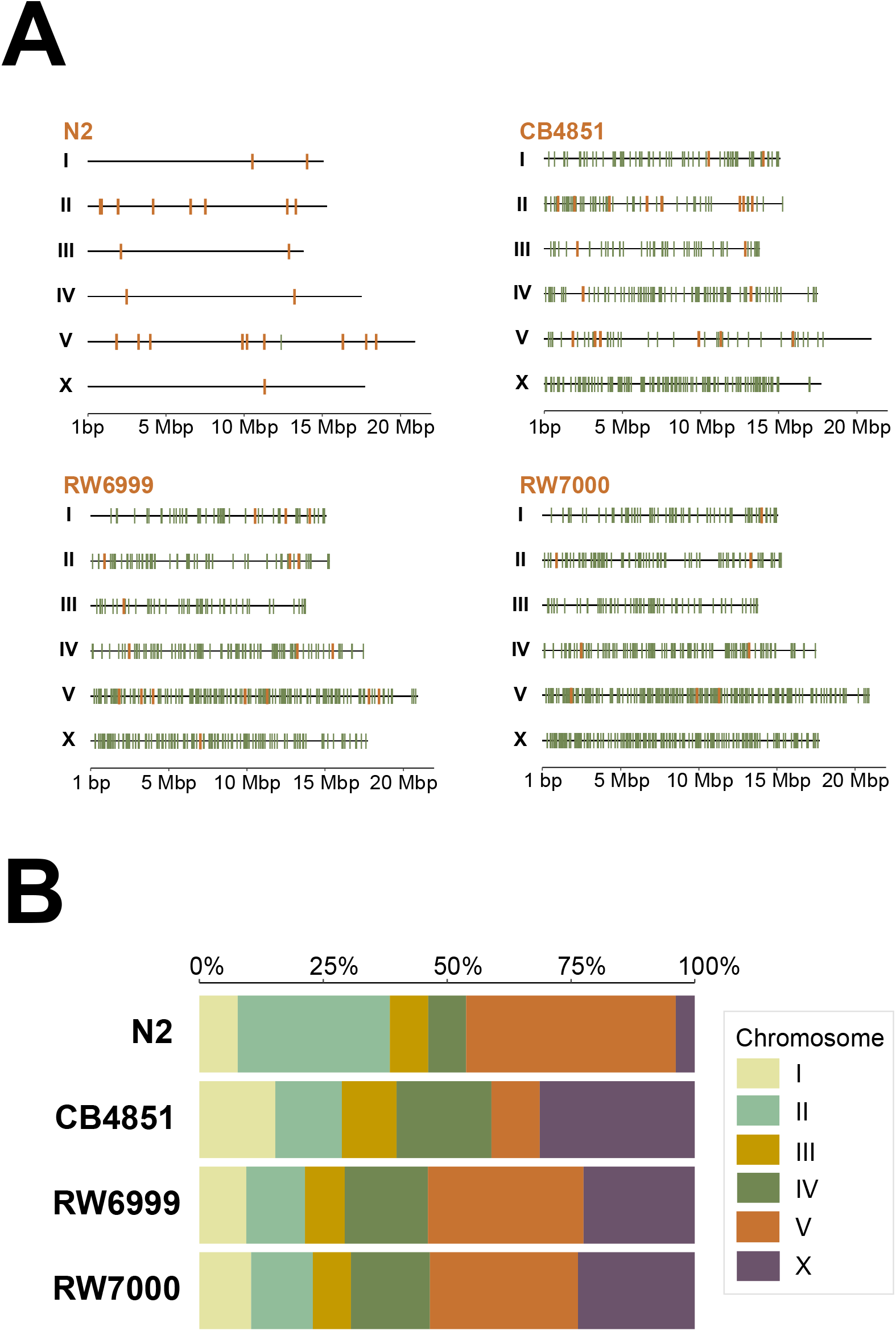
The chromosomal distribution of Tc*1* elements in the Bergerac strains relative to N2. (A) Genomic map of Tc*1* insertions within each strain. Non-reference Tc*1* insertions are displayed in green, while reference Tc*1* sites in Bristol N2 are shown in orange. (B) The proportion of Tc*1* elements by chromosome. Tc*1* elements were significantly overrepresented on the X chromosome in CB4851 (*χ*^2^ = 51.87, *p* = 5.92 × 10^-13^), and chromosomes V and X in RW6999 (*χ*^2^ = 49.06, *p* = 2.16 × 10^-9^) and RW7000 (*χ*^2^ = 50.88, *p* = 9.15 × 10^-10^).

The genome was additionally split into arms, cores, and tips based on recombination domains designated by Rockman and Kruglyak (2009) (**Figure 7**). Gene-poor high-recombination arms, gene-rich low-recombination cores, and gene-poor low-recombination tips comprise 45.7%, 47% and 7.3% of the *C. elegans* genome, respectively. The distribution of Tc*1* insertions in arms and cores was not significantly different from random expectation within N2 (*χ*^2^ = 2.66, *p* = 0.21), CB4851 (*χ*^2^ = 0.47, *p* = 0.49), RW6999 (*χ*^2^ = 4.39, *p* = 0.11) and RW7000 (*χ*^2^ = 4.81, *p* = 0.11) corrected for multiple-comparisons (Holm-Bonferroni method; Holm 1979). Furthermore, there was no significant difference between cores and arms in Tc*1* insertions that are shared between Bergerac strains and insertions that are unique to a particular strain (**Supplemental Figure S2**; *G* = 3.24, *p* = 0.072). Additional analysis of chromosomal tips was excluded because they represent such a small proportion of the genome.

**Figure 7.**
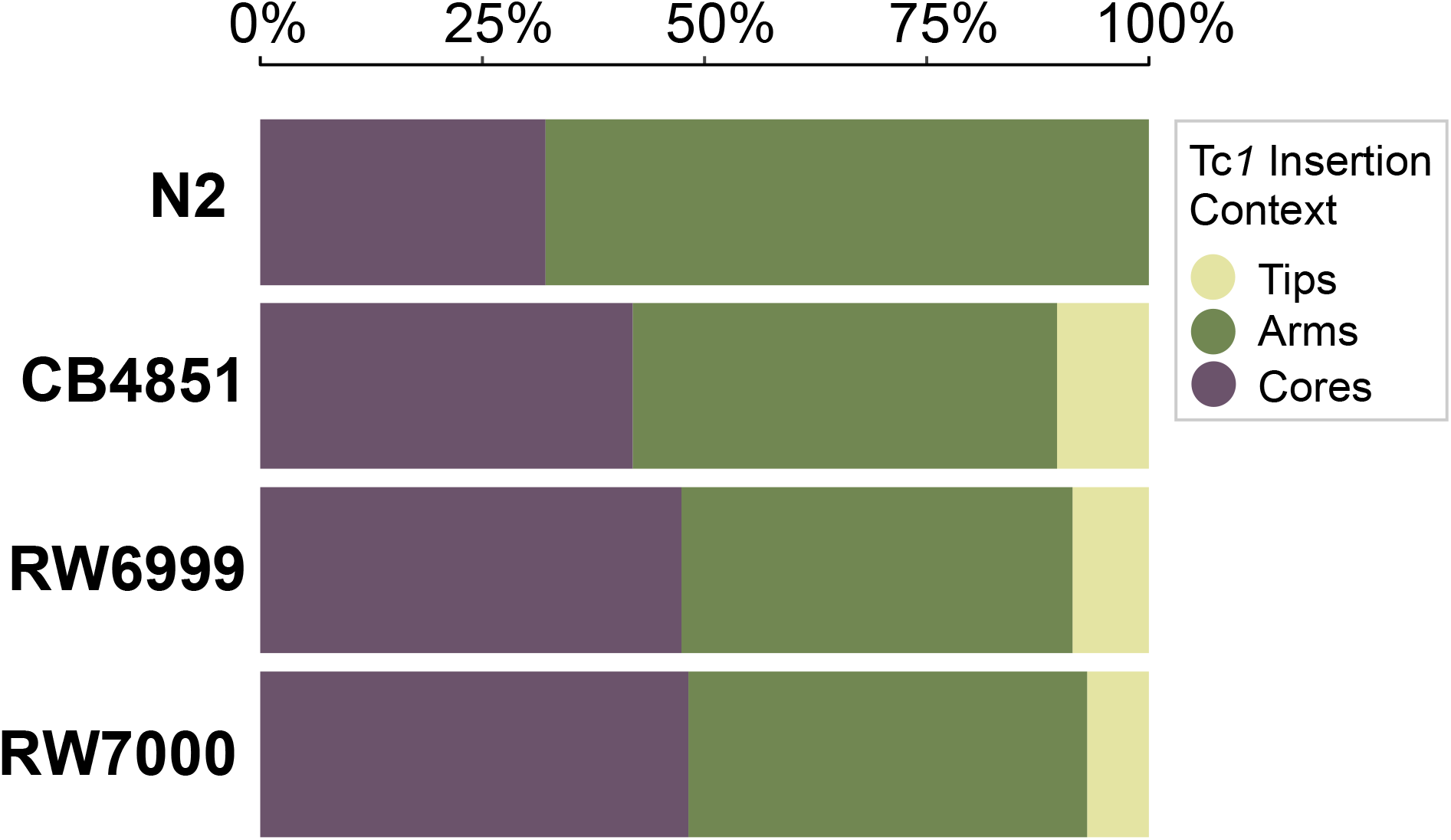
The proportions of Tc*1* elements located in arms, cores and tips. Arms are associated with high recombination rates whereas cores and tips have low recombination rates. There were no significant differences in the distribution of Tc*1* between these three domains (N2: *χ*^2^ = 2.66, *p* = 0.21; CB4851: *χ*^2^ = 0.47, *p* = 0.49; RW6999: *χ*^2^ = 4.39, *p* = 0.11; RW7000: *χ*^2^ = 4.81, *p* = 0.11).

To ascertain transposition into different chromatin environments, five previously identified broad patterns of histone modification (Liu *et al*. 2011) were used to gate Tc*1* insertions in the three Bergerac strains relative to the Bristol N2 strain. These five patterns of histone modification are as follows: (i) lowly expressed genes (H3K27me3), (ii) repetitive regions (H3K9me1/2/3), (iii) dosage compensation (H3K27me1, H4K20me1), (iv) promoters of highly expressed genes (H3K4me1/2/3, H3K27ac, H4K8ac, H4K16ac), and (v) highly expressed genes (H3K36me3, H3K79me1/2/3). As there was potential overlap between histone modification context, each category was tested individually for each strain. Interestingly, all Tc*1* locations identified in N2, both in the reference genome and in our laboratory isolate, inhabited regions associated with H3K27me3 or H3K9me1/2/3, and canonically identified as repressed in the *C. elegans* genome (**Figures 8A and 8B**), while being absent from regions associated with high expression (**Figures 8C and 8D**).

**Figure 8.**
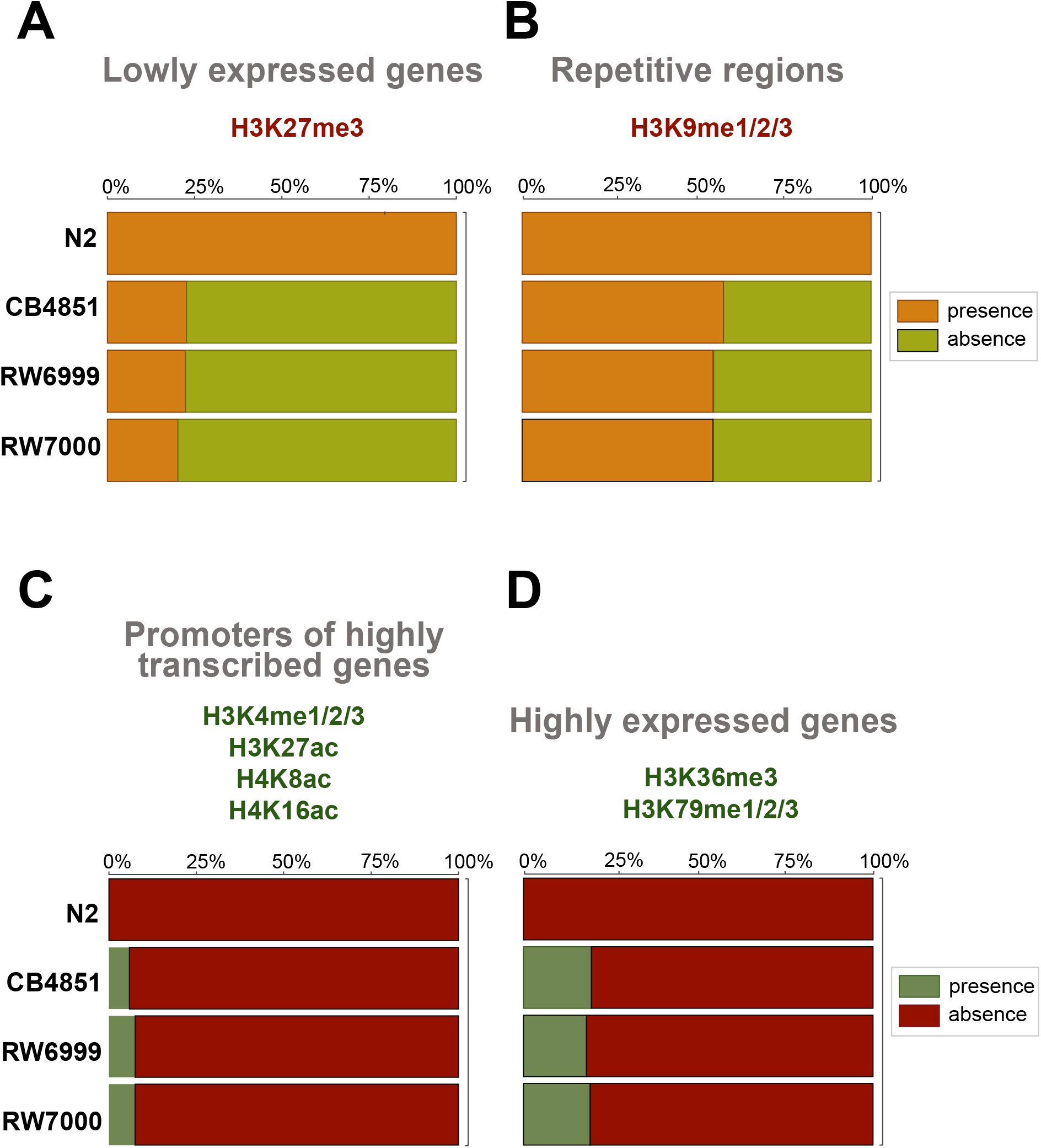
The proportions of Tc*1* insertions across different chromatin domains. (A) Histone modification H3K27me3 is associated with repressed chromatin and lowly expressed genes. All of the Tc*1* positions in N2 were found in this domain. Tc*1* insertions were more abundant in regions associated with H3K27me3 than expected by chance in CB4851 (*χ*^2^ = 7.44, *p* = 0.013) and RW6999 (*χ*^2^ = 8.82, *p* = 0.0089), but not RW7000 (*χ*^2^ = 2.82, *p* = 0.09). (B) Histone modifications H3K9mme1/2/3 are associated with repetitive DNA. All of the Tc*1* positions in N2 were found in these domains. Tc*1* insertions were overrepresented in H3K9me1/2/3 domains in strains RW6999 (*χ*^2^ = 5.06, *p* = 0.049) and RW7000 (*χ*^2^ = 7.06, *p* = 0.024), but not in CB4851 (*χ*^2^ = 7.06, *p* = 0.44). (C) Histone modifications H3K4me1/2/3, H3K27ac, H4K8ac, H4K16ac are associated with promoters of highly expressed genes. In N2, Tc*1* insertions were entirely absent from these domains. Tc*1* insertions were underrepresented in domains associated with H3K4me1/2/3, H3K27ac, H4K8ac, H4K16ac and promoters of in all three Bergerac strains; CB4851 (*χ*^2^ = 10.91, *p* = 0.00287), RW6999 (*χ*^2^ = 5.96, *p* = 0.0184); RW7000 (*χ*^2^ = 6.79, *p* = 0.0184). (D) Histone modifications H3K36me3 and H3K79me1/2/3 are associated with highly expressed genes. In N2, Tc*1* insertions were entirely absent from these domains. In all three Bergerac strains, Tc*1* insertions were underrepresented in these domains (CB4851: *χ*^2^ = 48.33, *p* = 3.60 × 10^-12^; RW6999: *χ*^2^ = 75.28, *p* = 8.17 × 10^-18^; RW7000: *χ*^2^ = 82.71, *p* = 2.85 × 10^-19^).

**Figure 9.**
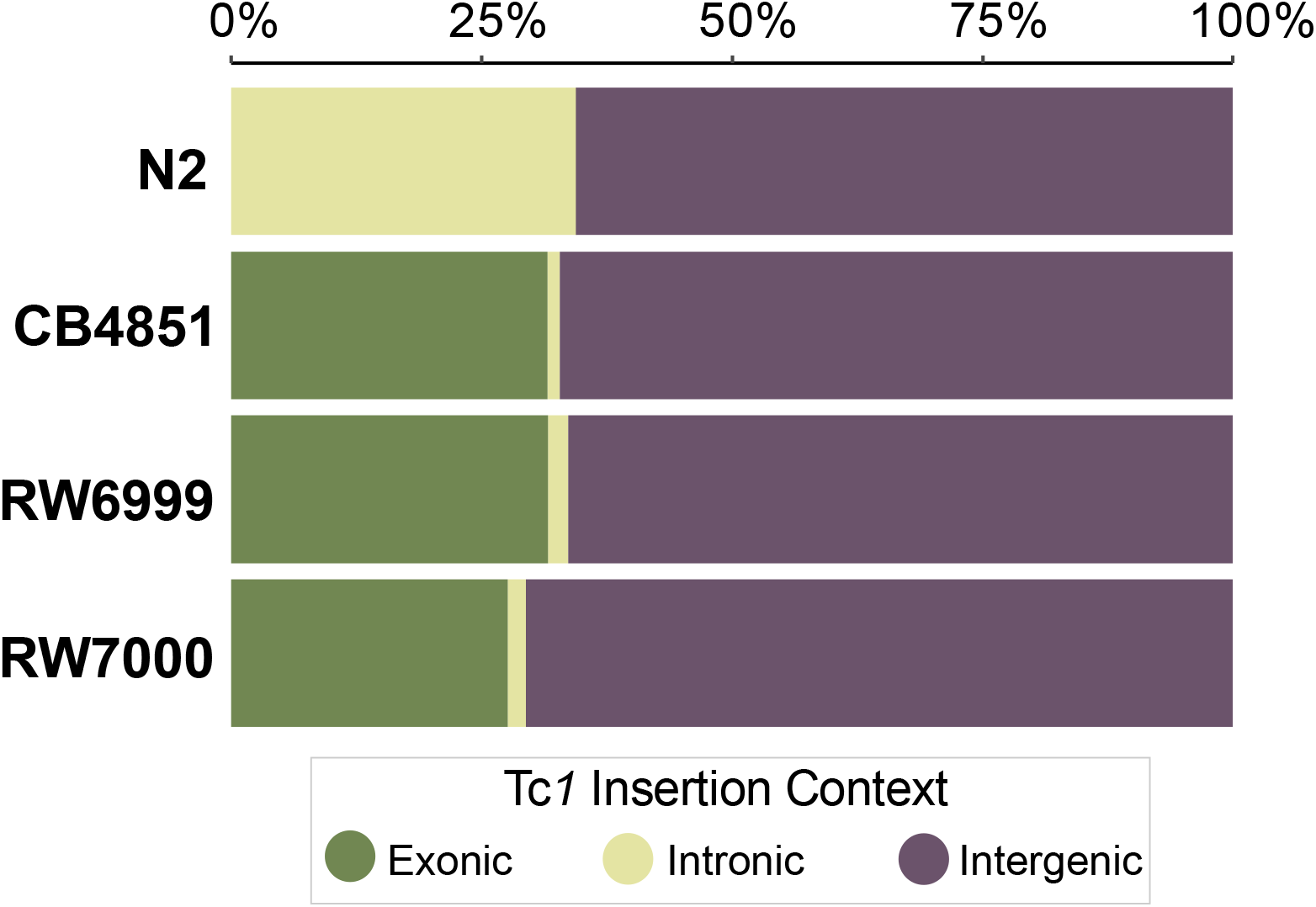
The proportion of Tc*1* elements in exons, introns, and intergenic regions of Bergerac strains in comparison to Bristol N2. In N2, the Tc*1* insertions were only found in introns and intergenic regions. The proportion of Tc*1* insertions in exons, introns and intergenic regions was significantly different between Bristol N2 and the Bergerac strains (*G* = 54.26, *p* = 6.54 × 10^-10^, *df* = 6), with the Bergerac strains showing a far larger and fewer proportion of Tc*1* insertions in exonic regions and intronic regions, respectively. There was no significant difference among the three Bergerac strains in the proportion of Tc*1* insertions in exons, introns, and intergenic regions (*G* = 4.28, *p* = 0.37, *df* = 4).

No significant preferences were found in Tc*1* insertions in domains associated with dosage compensation and enriched on the X chromosome (H3K27me1, H4K20me1) in any of the Bergerac strains (CB4851: *χ*^2^ = 0.003, *p* = 1; RW6999: *χ*^2^ = 2.12, *p* = 0.40; RW7000: *χ*^2^ = 0.44, *p* = 1, Holm-Bonferroni corrected) (not shown in **Figure 8**). There were no significant differences between the three Bergerac strains with regards to Tc*1* insertions in any of the five chromatin modification regions analyzed here (four shown in **Figure 8**). In heterochromatic regions associated with H3K27me3 and low gene expression (**Figure 8A**), Tc*1* insertions were more abundant than expected by chance in CB4851 (*χ*^2^ = 7.44, *p* = 0.013) and RW6999 (*χ*^2^ = 8.82, *p* = 0.0089), but not RW7000 (*χ*^2^ = 2.82, *p* = 0.09) after correcting for multiple comparisons (Holm 1979). Chromatin regions associated with H3K9me1/2/3, repetitive DNA and low gene expression (**Figure 8B**), were also slightly enriched for Tc*1* insertions in RW6999 (*χ*^2^ = 5.06, *p* = 0.049) and RW7000 (*χ*^2^ = 7.06, *p* = 0.024), but not in CB4851 (*χ*^2^ = 7.06, *p* = 0.44). Domains associated with H3K36me3, H3K79me1/2/3 and high gene expression (**Figure 8D**) had fewer Tc*1* insertions than expected by chance in all three Bergerac strains (CB4851: *χ*^2^ = 48.33, *p* = 3.60 × 10^-12^; RW6999: *χ*^2^ = 75.28, *p* = 8.17 × 10^-18^; RW7000: *χ*^2^ = 82.71, *p* = 2.85 × 10^-19^, Holm-Bonferroni corrected). Similarly, domains associated with promoters of highly expressed genes (H3K4me1/2/3, H3K27ac, H4K8ac, H4K16ac, **Figure 8C**) had fewer Tc*1* insertions than expected by chance (CB4851: *χ*^2^ = 10.91, *p* = 0.00287; RW6999: *χ*^2^ = 5.96, *p* = 0.0184; RW7000: *χ*^2^ = 6.79, *p* = 0.0184, Holm-Bonferroni corrected). Some of the Tc*1* insertions associated with highly expressed genes in the Bergerac strains were found in germline enriched genes (Han *et al*. 2019). CB4851, RW6999, and RW7000 harbored 10, 17, and 21 Tc*1* insertions within or near highly expressed germline enriched genes, respectively. In sum, repressed chromatin had excess insertions and transcriptionally active chromatin had fewer Tc*1* insertions than expected by chance. However, Tc*1* insertions are significantly more likely in transcriptionally active chromatin in the Bergerac strains than in N2 (Fisher’s exact test: N2 *vs.* CB4851, *p_adj_* = 0.0243; N2 *vs.* RW6999, *p_adj_* = 0.0272; N2 *vs.* RW7000, *p_adj_* = 0.0243).

There were significant differences between the four strains in the distribution of Tc*1* elements in exons, introns, and intergenic regions (*G* = 54.26, *p* = 6.5 × 10^-10^). None of the Tc*1* insertions in N2 were present in exons; rather they appear to be overrepresented in introns (Fischer’s exact: *p* = 1.3 × 10^-5^). The distribution of Tc*1* is not significantly different across exons, introns, and intergenic regions between the three Bergerac strains (*G* = 4.28, *p* = 0.28). However, Tc*1* appears to be underrepresented in exons and overrepresented in introns and intergenic regions in RW6999 (*χ*^2^ = 10.20, *p* = 0.012, adjusted for multiple comparison) and RW7000 (*χ*^2^ = 22.32, *p* = 5.6 × 10^-5^, *p* adjusted by Holm-Bonferroni correction). The underrepresentation of Tc*1* in exons could be due to selection. However, potential Tc*1* insertion sites are also more abundant in introns and intergenic regions due in part to their higher A/T content compared to exons. If we recalculate the expected number of Tc*1* insertions in exons, introns and intergenic regions based on the frequency of the consensus integration sequence in these categories, Tc*1* insertions are overrepresented in exons and underrepresented in intergenic regions (CB4851: *χ*^2^ = 38.61, *p* = 8.27 × 10^-9^; RW6999: *χ*^2^ = 59.53, *p* = 3.55 × 10^-13^; RW7000: *χ*^2^ = 35.14, *p* = 2.35 × 10^-8^, *p* adjusted by Holm-Bonferroni correction). For example, in Bergerac strain RW6999, the expected number of Tc*1* insertions in exons is 121 based on the frequency of consensus insertions sites whereas the observed number is 190.

### Mutations in the Bergerac strains

*Tc1* insertions were found to disrupt the exons of many protein-coding genes. Since the insertion of TEs into exons can lead to a disruption or total knock-out of gene function (Orgel and Crick 1980; Muñoz-López and García-Pérez 2010), the RNAi phenotypes for these genes provides a prediction for the effect of these insertions. These RNAi phenotypes (**Supplemental Table S10**) include many phenotypes observed in the phenotypic assays, including reduced brood size, slow growth, embryonic lethality, shortened life span, and locomotion defects. In addition, the Bergerac strains share 6,191 homozygous SNPs/indels after filtering (see Materials and Methods). These mutations were predicted to result in an amino acid change in 384 genes (**Supplemental Table S11**) and 122 of these mutations were predicted to have a deleterious effect on protein function by SIFT (Vaser *et al*. 2016). No candidate mutations were identified in genes known to be relevant for the small RNA pathways linked to regulation of TE activity (**Supplemental Table S11**; Billi *et al*. 2014). However, the protein *abt-1* was annotated by WormBase as being involved in germline RNAi. Despite a lack of damage to sRNA-related genes, several genes related to meiosis and recombination (*him-8*, *msh-5*, *rec-8*, *zyg-12*, and *dsb-1*) were predicted to have deleterious amino acid changes by SIFT (Vaser *et al*. 2016). The MiModd package also produced calls for deletions in the Bergerac strains. However, no exonic deletions in genes potentially involved in TE suppression were identified.

### Dating the Divergence of Bergerac and N2 (Bristol) strains of C. elegans

In order to generate divergence time estimates among the Bergerac strains and the N2 (Bristol) strain, we employed WGS data to determine mtDNA genetic divergence (SNPs) between N2 and each of the three focal Bergerac strains. N2−RW7000, N2−RW6999, and N2−CB4851 differed by four SNPs each. We assumed (i) a spontaneous base substitution rate of 4.32 × 10^-8^ /site/generation for the *C. elegans* mtDNA genome (Konrad *et al*. 2017), and (ii) three estimates of generation time (14-, 30- and 60-day generation) in wild *C. elegans* which is more apt based on the ecology of the species (Cutter 2008). Assuming a generation time of 14, 30, and 60 days, wild *C. elegans* are expected to have approximately 26, 12, and six generations per year, respectively. Under this range of generation time, all three Bergerac strains are estimated to have diverged from N2 between 129−552 years (14-day generation: 129 years; 30-day generation: 276 years; 60-day generation: 552 years). Finally, to place the Bergerac strains in a phylogenetic context, we ran a maximum likelihood analysis (PhyML) on 11 strains, which included the three Bergerac strains and N2 using both mtDNA and nuclear SNPs. The phylogenetic tree matched the expected relationships, with the French natural isolates JU394 and JU2565 displaying the closest relationship to the Bergerac strains, and the Bergerac strains forming a monophyletic group on the tree (**Supplemental Figure S2**) (Cook *et al*. 2017).

## DISCUSSION

The Bergerac and the N2 (Bristol) strains were the first two laboratory isolates of *C. elegans* and hence, of historical importance in the development of the species as a model organism (Riddle *et al*. 1997). Laboratory culturing of the Bergerac strain preceded that of N2 by several decades (Nigon 1949; Nigon and Dogherty 1949). However, the Bergerac strain displayed temperature-sensitivity (Fatt and Dogherty 1963), low fertility (Abdulkader and Brun 1980; Lee *et al*. 2016), and other phenotypic abnormalities including uncoordinated movement (Emmons *et al*. 1983; Hodgkin and Doniach 1997) which ultimately led to its supplantation in the laboratory by the more phenotypically robust Bristol N2 strain (Brenner 1974; Sulston and Brenner 1974). A series of well-executed studies in succession over a decade from the late 1970s to late 1980s served to elucidate the genetic bases of these phenotypic peculiarities that have since become characteristic of the original Bergerac isolate and its descendants (see Introduction), namely the proliferation of a unique DNA transposon called Tc*1* (Emmons *et al*. 1979, 1983; Liao *et al*. 1983; Eide and Anderson 1985; Egilmez *et al*. 1995).

Sequencing of a Tc*1* element showed it to be 1,610 bp in length (coding potential of 343 aa) with short inverted terminal repeats of 54 bp (Rosenzweig *et al*. 1983a). Interestingly, all copies of Tc*1* in *C. elegans* appeared to be uniform in length and largely conserved in their DNA sequence, suggesting that Tc*1* coded for products mediating its own transcription (Liao *et al*. 1983) or that the regulatory mechanism for transposition required a full-length element (Mori *et al*. 1988). The transposition of Tc*1* was shown to be strain-specific (Eide and Anderson 1985) with the Bergerac strain BO (RW7000) capable of generating spontaneous mutations in the *unc-22* locus at a per generation rate of approximately 10^-4^, and three orders of magnitude higher than the baseline spontaneous mutation rate at the same locus in the N2 genetic background (Moerman and Waterston 1984). Furthermore, whilst both the Bergerac (RW7000 subclone) and N2 strains exhibited high frequencies of somatic excision (Emmons and Yesner 1984; Emmons *et al*. 1986), the Bergerac strain was unique in having active germline transposition (Collins *et al*. 1987). It should be mentioned that the history and nomenclature of the Bergerac strains can be confusing given that clones were shared with multiple laboratories over a period of decades and sublineages were often given new names (reviewed in Nigon and Félix 2017). While RW6999 is regarded as a subclone of Bergerac RW7000/BO/LY, the Bergerac strain CB4851 has high Tc*1* content (Hodgkin and Doniach 1997) but has been reported to lack the active germline transposition of Tc*1* (Nigon and Félix 2017) as observed in the RW7000 and RW6999 strains (Moerman and Waterston 1984; Eide and Anderson 1985). Despite a decade of sustained focus in the pre-genomics era, a modern genomic analysis of the genomic distribution of TEs, a precise quantification of their relative fitness, and further investigations into a possible cause for TE deregulation in the Bergerac strains is hitherto lacking.

The transposable element Tc*1* was shown to be unique to *C. elegans* and while present in low copy-number in the majority of *C. elegans* strains examined, the Bergerac strain and its descendants were classified as high copy-number strains. Emmons *et al*. (1983) were the first to use Southern blotting to estimate genomic Tc*1* copy-number in the N2 and Bergerac strain RW7000 (also referred to as the BO subclone) as 20 ± 5 and 200 ± 50, respectively. Liao *et al*. (1983) estimated Tc*1* copy-number in the N2 and Bergerac strain RW7000 (also referred to as the LY subclone) to be 25-30 and several hundred copies, respectively. Eide and Anderson (1985) mention Bergerac RW7000/BO/LY lineage as possessing 250 Tc*1* copies as per Emmons *et al*. (1983) and Liao *et al*. (1983). Mori *et al*. (1988) subsequently found the RW7000/BO/LY lineage to harbor >300 Tc*1* copies. Hence, Southern blotting techniques established a range of 25-30 and 200 to >300 Tc*1* copies for the N2 and RW7000 strain, respectively. Recognizing the limitation of Southern blots in the accurate estimation of Tc*1* in high-copy strains, Egilmez *et al*. (1995) opted to use DNA dot-blots. N2 and Bergerac RW7000/BO/LY were estimated to possess 31 and 515 ± 48 Tc*1* copies, respectively (Egilmez *et al*. 1995). More recently, Larichhia *et al*. (2017) were the first to estimate transposon abundance in several hundred *C. elegans* isolates from Illumina pair-end sequencing data. One Bergerac sublineage, CB4851, was included in their study and estimated to possess 406 Tc*1* copies (Laricchia *et al*. 2017). However, because there are no existing previous hybridization-based estimates of Tc*1* count for CB4851, it is not possible to compare transposon number estimates generated from hybridization versus WGS data. This study offers new estimates of Tc*1* counts for three Bergerac strains based on both (i) bioinformatic analyses of WGS data, and (ii) independent confirmation by ddPCR. We determined far higher copy-number estimates of Tc*1* in the Bergerac strains than preceding studies with median values of 451, 581 and 748 copies for CB4851, RW6999, and RW7000, respectively. In the best-studied RW7000 Bergerac strain, our estimate of Tc*1* copy-number is more than 2× greater than past studies relying on Southern hybridizations (Emmons *et al*. 1983; Liao *et al*. 1983; Mori *et al*. 1988) and approximately 1.5× greater than an estimate based on DNA dot-blots (Egilmez *et al*. 1995). The discrepancy in our estimates of Tc*1* copy-number and those of past studies relying on Southern hybridizations and DNA dot-blots is likely due to drawbacks associated with the latter techniques, namely membrane saturation limiting resolution of copy-number when counts exceed ∼100. The five bioinformatic methods employed to estimate TE copy-number were consistent with each other, both in the number of calls per strain and in the differences in Tc*1* copy-number between the strains. Furthermore, 90% and 70% of the calls were shared by more than one method and by all methods, respectively. Nevertheless, these median numbers of Tc*1* are likely to be underestimates as both sequence read coverage and ddPCR yielded even higher estimates of Tc*1* copy number in the three Bergerac strains.

Both our ddPCR and bioinformatic approaches demonstrate that Tc*1* copy-number can vary considerably among the three Bergerac sublineages/strains examined in this study (median values of 451, 581 and 748 copies for CB4851, RW6999, and RW7000, respectively). This among-strain variation in Tc*1* copy-number could be owing to either one or a combination of at least two factors. First, it is possible that some Bergerac sublineages have evolved a mechanism of TE regulation that limits further proliferation of Tc*1*. For example, CB4851 is thought to lack the active germline transposition characteristic of the RW7000 and RW6999 strains (Nigon and Félix 2017). Second, the complete history of laboratory propagation for these strains over multiple decades is obscure, with some strains possibly having been subjected to lengthier periods of cryopreservation or laboratory evolution than others.

Aside from Tc*1*, the only consistent difference in TE copy-number between the Bergerac strains and N2 was for Tc*2*. The average number of Tc*2* elements was 20 and three in the Bergerac strains and N2, respectively, and normalized read depth suggests 25−30 copies of Tc*2* in the Bergerac strains and four in N2. Tc*2*, originally discovered in a Bergerac strain, was upon its discovery found in higher copy-number Tc*1* strains than in low-copy number Tc*1* strains (Levitt and Emmons 1989). This coincidence suggested that the proliferation of Tc*1* and Tc*2* was due to shared copy-number regulation mechanisms (Levitt and Emmons 1989). Since then, many genes have been discovered in *C. elegans* that influence the transposition of multiple TEs (Vastenhouw *et al*. 2003; Lee *et al*. 2012; Weick and Miska 2014; McMurchy *et al*. 2017; Wallis *et al*. 2019). In contrast to the striking differences among the Bergerac strains with respect to the number of Tc*1* insertions, Tc*2* copy-number and locations are remarkably well-conserved. It appears that the explosive proliferation and divergence of Tc*1* insertions in the Bergerac strains is caused by factors specific to Tc*1* and not shared with other TEs in *C. elegans*.

While Bergerac strains have long been observed to harbor mutational defects (Wood *et al*. 1980; Emmons *et al*. 1983), this study is the first to offer precise quantification of their low composite fitness via phenotypic assays of four fitness-related traits (productivity, survivorship to adulthood, developmental rate, and longevity). Notably, all three Bergerac strains differ significantly from N2 with respect to each of the four fitness traits; in each case, their mean fitness trait values are far lower than N2. Productivity and survivorship to adulthood are two traits considered most germane to organismal fitness. The three Bergerac strains exhibited an extensive reduction (65-86%) in productivity relative to wild type N2. To place this in perspective, spontaneous mutation accumulation (MA) lines of *C. elegans* subjected to extreme genetic drift and inbreeding for >400 consecutive generations displayed an average 44-55% reduction in productivity (Katju *et al*. 2015, 2018). Mean survivorship to adulthood in the three Bergerac strains was reduced by 7-15% relative to wild type N2. In contrast to productivity, this reduction in survivorship of the Bergerac strains is on par with the 12-19% decline observed in the same long-term spontaneous MA lines of *C. elegans* evolved over 400 generations (Katju *et al*. 2015, 2018). The Bergerac strains also exhibited reduced longevity (29-36% lower relative to N2) as well as delayed developmental rate (9-20% longer than N2). These experiments do not bear on the question whether TE proliferation alone directly contributes to all observable fitness decline in the Bergerac strains. Indeed, an independent mutation in the *zyg-12* gene in the Bergerac strain renders it almost completely sterile at 25°C (Malone *et al*. 2003). However, TE mobilization events have been shown to have a direct negative impact on host fitness; notable examples include the reduction of host fertility and viability following *P*-element invasion in *Drosophila* (Kidwell and Novy 1979), decreased fitness and egg hatchability in laboratory lines of *D. melanogaster* with elevated TE transposition rates (Pasyukova *et al*. 2004), and fitness loss in *Drosophila* mutation accumulation lines following *copia* insertions (Houle and Nuzhdin 2004). It is therefore reasonable to assume that the low fitness of the Bergerac strain and its derivatives is owing largely to the unchecked proliferation of TEs.

In addition to confirming that all Bergerac strains had significantly lower fitness than N2, our study also quantified significant inter-strain differences among the three Bergerac strains studied here with respect to fitness. RW7000 (or Bergerac-BO), used in many transposon-tagging studies in the 1990s (Korswagen *et al*. 1996), exhibited the lowest means for three traits (productivity, and survivorship, and longevity). RW7000 also possesses the highest Tc*1* copy-number among the three strains studied here. Interestingly, the strain RW6999, listed as an RW subclone of RW7000 by the Caenorhabditis Genetics Center (CGC), displayed the highest fitness among the Bergerac strains, despite having the second-highest number of Tc*1* insertions. The hypothesis that TE proliferation is the sole cause of fitness differences between the three Bergerac strains is ruled out by these results. It appears that fitness is not simply correlated with copy-number of active Tc*1* elements but other determinants as well such as site(s) of genomic integration, and/or certain key loci with high mutator activity.

Bergerac strains are also known to display abnormal phenotypes with respect to locomotion, and are often referred to as “uncoordinated” (Brenner 1974; Hodgkin and Doniach 1997; Emmons *et al*. 2003; Nigon and Félix 2017). The *unc-22* gene on Chr. IV plays an important role in muscle structure and function (Moerman *et al*. 1988), and is a frequent target for Tc*1* element insertions in the Bergerac strains (Moerman *et al*. 1986). As was the case with fitness-related traits, all three Bergerac strains had significantly reduced mean body length (16-20% reduction) and body area (17-31%) relative to N2. In addition, some Bergerac strains exhibit significant deviations from N2 with respect to locomotion. RW6999 and RW7000 have significant reductions in speed (32% and 70%, respectively) relative to N2. In addition, RW7000 worms displayed a compromised ability to move forward and turn, resulting in a significant increase in direction change (113%) relative to N2. Hence, it appears that RW7000 worms have the most severe impairment in locomotion relative to N2. It is likely that Tc*1* insertions have contributed to the observed changes in these quantitative traits in conjunction with other independent mutations in the unique evolutionary trajectory of the Bergerac ancestor and its derived lineages. One mutation present in all three Bergerac strains was a nonsynonymous base substitution in an olfactory G-protein-coupled receptor (GPCR) *str-208* yielding a Cys226Ser/Tyr. This mutation may result in improper folding of the protein, with possible inhibitory effects on proper signaling and sensation of external stimuli. The inability to properly relay/convey external stimuli could explain the erratic movement exhibited by the Bergerac strains in the motility assays. The Bergerac strains were additionally noted to leave the area of the bacterial feeding lawn or position themselves around the bacterial seed instead of entering to feed. This behavior could be rooted in a dysfunctional olfactory system and possibly contribute to diminished fitness.

The genomes of multicellular eukaryotes are an assemblage of DNA sequences necessary for function as well as repetitive DNA sequences that may not contribute to organismal function. This latter component, which may constitute a large fraction of eukaryotic genomes, includes TEs that can be detrimental to the host genome when they transpose into coding regions thereby causing gene disruption. The genome has therefore been likened to an ecological community with TEs being functionally regarded as parasitic invaders with the host genome under selective pressure to actively engage in tackling this TE spread (Brookfield 2005; Bourque *et al*. 2018). Hence, the distribution and abundance of TEs across the genome is frequently nonrandom due to a combination of natural selection, as well as TE integration site preferences, and deletions (Bourque *et al*. 2018). TEs are a source of both germline and somatic mutations, contribute to ectopic recombination and can interfere with normal gene expression, all of which are, on average, deleterious to host fitness. Consequently, TEs are observed less frequently within coding and regulatory DNA sequences than expected by chance. Furthermore, the distribution of TEs in genomes can also depend on the local recombination rate due to the effects of natural selection against both ectopic exchange and insertional mutations. For example, the numbers of DNA-based transposable elements are negatively correlated with recombination rate in the *Drosophila melanogaster* genome (Rizzon *et al*. 2002). Assuming that opportunities for ectopic recombination between TEs increase with the local recombination frequency, the presence of TEs is more likely to be detrimental in regions with high recombination rates compared to low recombination rates (Langley *et al*. 1988). Moreover, natural selection is less efficient in regions with low recombination relative to high recombination which predicts that deleterious TE insertions are more likely to accumulate in regions of low recombination (Hill and Robertson, 1966; Dolgin and Charlesworth 2008). In addition, insertional preferences may vary with non-random distribution of features such as suitable integration motifs, recombination rate, and chromatin organization (Sultana *et al*. 2017). Tc*1* transpositions in *C. elegans* have been shown to occur more frequently both (i) within than between chromosomes, and (ii) to proximate than distant locations on the same chromosome (Fischer *et al*. 2003).

Consequently, this bias towards proximate locations violates the usual assumption of independence in statistical tests. If, for now, we ignore these limitations to interpreting the genome-wide distribution of Tc*1* elements, we find that Tc*1* is overrepresented on Chr. V in strains RW6999 and RW7000, and on the X chromosome in all three Bergerac strains after correcting for chromosome length. In *C. elegans*, the chromosome arms are relatively gene-poor regions but have higher recombination frequency than chromosome cores, which are also gene-rich (Rockman and Kruglyak, 2009). In contrast to the relationship between recombination rate and TEs in *Drosophila*, TE abundance in *C. elegans* is greater in the high-recombination arms than in the low-recombination cores (Duret *et al*. 2000; Laricchia *et al*. 2017). We did not find a significant difference in the abundance of Tc*1* elements between the chromosome arms and cores across the three Bergerac strains analyzed here. In this regard, our results are similar to an earlier analysis of TEs in *C. elegans* that also failed to find a significant relationship between recombination rate and Tc*1* copy-number (Duret *et al*. 2000).

Tc*1* is underrepresented in exons which is consistent with selection against strongly deleterious insertions in coding regions of the genome. However, the consensus Tc*1* integration motif is more commonly found in introns and intergenic regions. The underrepresentation of Tc*1* abundance in exons could therefore primarily be a function of the availability of suitable integration motifs. In a similar vein, Tc*1* insertions in the Bergerac strains were slightly less and more abundant than expected by chance in transcriptionally active and repressed chromatin, respectively. This pattern could arise by insertional preference into repressed chromatin, unequal distribution of integration motifs between repressed and transcriptionally active chromatin, or selection against insertions into transcriptionally active chromatin. Indeed, there is strong purifying selection on TE expression in experimental populations of *C. elegans* (Bergthorsson *et al*. 2020).

Furthermore, there is a striking difference between the Bristol N2 and the Bergerac strains in that the former sequenced for this study retain Tc*1* insertions exclusively in repressive chromatin. The Bergerac strains contained none of the alleles that have previously been found to derepress Tc*1* activity (Vastenhouw *et al*. 2003; Lee *et al*. 2012; Weick and Miska 2014; McMurchy *et al*. 2017; Wallis *et al*. 2019) and mapping putative mutator loci for Tc*1* in these strains suggested that the mutator activity was associated with multiple sites in the genome which were themselves mobile (Moerman and Waterston 1984; Mori *et al*. 1988). The presence of numerous Tc*1* elements insertions in transcriptionally active chromatin in the Bergerac strains could explain their high rate of germline transposition. Even if Tc*1* preferentially integrates into repressed chromatin, thus evading purifying selection to some degree, serendipitous insertions of Tc*1* into an active chromatin environment could create a proliferative loop of Tc*1* expansion which selection is simply unable to completely restrain.

The evolutionary history of the Bergerac strain and its derivatives remains obscure. The current Bergerac strain was isolated by Victor Nigon from his garden soil in Bergerac, France in 1944 (Nigon and Félix 2017). How evolutionary diverged are the Bergerac and the N2 strains? Both earlier and recent studies of *C. elegans* intraspecific genetic diversity have similarly observed that single-nucleotide variant (SNV) diversity is often shared among many natural isolates and that the species is characterized by lower levels of genetic diversity relative to other obligately outcrossing as well as facultatively selfing species in the genus *Caenorhabditis* (Denver *et al*. 2003; Sivasundar and Hey 2003; Cutter 2006; Dey *et al*. 2013; Thomas *et al*. 2015). Based on mtDNA genetic divergence among the three Bergerac strains and N2 and assuming three different estimates of generation time in the wild, we estimated a very evolutionary recent N2-Bergerac divergence range of 129-552 years. As expected, the three focal Bergerac strains in this study form a monophyletic clade and are more closely related to two French natural isolates, providing evidence of some biogeographic structure.

Another pertinent outstanding question is whether the high activity of Tc*1* transposition in the Bergerac germline originated in the wild or occurred following laboratory domestication. Interestingly, the subclones/derivatives of Bergerac display variation with respect to active germline transposition of Tc*1* (Emmons *et al*. 1983; Liao *et al*. 1983; Moerman and Waterston 1984). For example, the BO/RW7000 sublineage has active germline transposition (Moerman and Waterston 1984), extremely elevated Tc*1* copy-number and drastically reduced fitness (latter two features confirmed in this study), whereas CB4851 has a high transposon content (though not as elevated as BO/RW7000 and RW6999) but is thought to lack active germline transposition of Tc*1* (Nigon and Félix 2017). What evolutionary scenario would explain the active germline transposition of Tc*1* in some Bergerac sublineages and not others? Under one scenario, the wild ancestor of all Bergerac strains possessed active germline transposition of Tc*1* which has been suppressed (selection) or lost (genetic drift) in some Bergerac derivatives following laboratory domestication. A second scenario to be considered is the activation of Tc*1* germline transposition after introduction to the laboratory, sometime in the period between 1944 and 1969 (see schematic of tentative history of the Bergerac strain in Nigon and Félix 2017) followed by subsequent suppression in different Bergerac lineages. It is conceivable that laboratory selection for novel mutations resulted in the preferential retention of sublineages of Bergerac that generated more mutations due to germline Tc*1* activity. The question of how Tc*1*proliferation in the Bergerac was initiated is still open.

## ACKNOWLEDGMENTS

This research was supported by start-up funds from the Department of Veterinary Integrative Biosciences, College of Veterinary Medicine and Biomedical Sciences at Texas A&M University to V.K. and U.B.

## LITERATURE CITED

Abdulkader, N., and J. Brun, 1980 A temperature-sensitive mutant of *Caenorhabditis elegans* var. Bergerac affecting morphological and embryonic development. Genetics 51: 81−92. https://doi.org/10.1007/bf00133506

Adrion, J. R., M. J. Song, D. R. Schrider, M. W. Hahn, and S. Schaack, 2017 Genome-wide estimates of transposable element insertion and deletion rates in *Drosophila melanogaster*. Genome Biol. Evol. 9: 1329−1340. https://doi.org/10.1093/gbe/evx050

Almeida, M. V., M. A. Andrade-Navarro, and R. F. Ketting, 2019 Function and evolution of nematode RNAi pathways. Noncoding RNA 5: 8. https://doi.org/10.3390/ncrna5010008

Ambros, V., R. C. Lee, A. Lavanway, P. T. Williams, and D. Jewell, 2003 MicroRNAs and other tiny endogenous RNAs in C. elegans. Curr. Biol. 13: 807−818. https://doi.org/10.1016/s0960-9822(03)00287-2

Angstman, N. B., H. Frank, and C. Schmitz, 2016 Advanced behavioral analyses show that the presence of food causes subtle changes in *C. elegans* movement. Front. Behav. Neurosci. 10: 1−10. https://doi.org/10.3389/fnbeh.2016.00060

Ashe, A., A. Sapetschnig, E.-M. Weick, J. Mitchell, M. P. Bagijn et al., 2012 piRNAs can trigger a multigenerational epigenetic memory in the germline of C*. elegans*. Cell 150: 88–99. https://doi.org/10.1016/j.cell.2012.06.018

Bagijn, M. P., L. D. Goldstein, A. Sapetschnig, E. M. Weick, S. Bouasker et al., 2012 Function, targets, and evolution of *Caenorhabditis elegans* piRNAs. Science 337: 574–578. https://doi.org/10.1126/science.1220952

Batista, P. J., J. G. Ruby, J. M. Claycomb, R. Chiang, N. Fahlgren et al., 2008 PRG-1 and 21U-RNAs interact to form the piRNA complex required for fertility in *C. elegans*. Mol. Cell 31: 67–78. https://doi.org/10.1016/j.molcel.2008.06.002

Beltran, T., V. Shahrezaei, V. Katju, and P. Sarkies, 2020 Epimutations driven by small RNAs arise frequently but most have limited duration in *Caenorhabditis elegans*. Nat. Ecol. Evol. 4: 1539–1548. https://doi.org/10.1038/s41559-020-01293-z

Bergthorsson, U., C. J. Sheeba, A. Konrad, T. Belicard, T. Beltran et al., 2020 Long-term experimental evolution reveals purifying selection on piRNA-mediated control of transposable element expression. BMC Biol. 18: 162. https://doi.org/10.1186/s12915-020-00897-y

Billi, A. C., S. E. Fischer, and J. K. Kim, 2014 Endogenous RNAi pathways in C. elegans. WormBook May 7: 1−49. https://doi.org/10.1895/wormbook.1.170.1

Blumenstiel, J. P., 2011 Evolutionary dynamics of transposable elements in a small RNA world. Trends Genet. 27: 23–31. https://doi.org/10.1016/j.tig.2010.10.003

Bourque, G., K. H. Burns, M. Gehring, V. Gorbunova, A. Seluanov et al., 2018 Ten things you should know about transposable elements. Genome Biol. 19: 199. https://doi.org/10.1186/s13059-018-1577-z

Brenner. S., 1974 The genetics of *Caenorhabditis elegans*. Genetics 77:71–94. https://doi.org/10.1093/genetics/77.1.71

Brookfield, J. F. Y., 2005 The ecology of the genome - mobile DNA elements and their hosts. Nat. Rev. Genet. 6: 128−136. https://doi.org/10.1038/nrg1524

Byerly, L., R. C. Cassada, and R. L. Russell, 1976 The life cycle of the nematode *Caenorhabditis elegans*: I. Wild-type growth and reproduction. Dev. Biol. 51: 23–33. https://doi.org/10.1016/0012-1606(76)90119-6

C. elegans Sequencing Consortium, 1998 Genome sequence of the nematode C. elegans: a platform for investigating biology. Science 282: 2012–2018. https://doi.org/10.1126/science.282.5396.2012

Chuong, E. B., N. C. Elde, and C. Feschotte, 2017 Regulatory activities of transposable elements: from conflicts to benefits. Nat. Rev. Genet. 18: 71–86. https://doi.org/10.1038/nrg.2016.139

Cingolani, P., A. Platts, L. L. Wang, M. Coon, T. Nguyen, et al., 2012 A program for annotating and predicting the effects of single nucleotide polymorphisms, SnpEff: SNPs in the genome of Drosophila melanogaster strain w1118; iso-2; iso-3. Fly 6: 80–92. https://doi.org/10.4161/fly.19695

Collins, J., B. Saari, and P. Anderson, 1987 Activation of a transposable element in the germ line but not the soma of *Caenorhabditis elegans*. Nature 328: 726–728. https://doi.org/10.1038/328726a0

Cook, D. E., S. Zdraljevic, J. P. Roberts, and E. C. Andersen, 2017 CeNDR, the *Caenorhabditis elegans* natural diversity resource. Nucleic Acids Res. 45: 650–657. https://doi.org/10.1093/nar/gkw893

Cutter, A. D., 2006 Nucleotide polymorphism and linkage disequilibrium in wild populations of the partial selfer *Caenorhabditis elegans*. Genetics 172:171–184. https://doi.org/10.1534/genetics.105.048207

Cutter, A. D., 2008 Divergence times in *Caenorhabditis* and *Drosophila* inferred from direct estimates of the neutral mutation rate. Mol. Biol. Evol. 25: 778–786. https://doi.org/10.1093/molbev/msn024

Das, P. P., M. P. Bagijn, L. D. Goldstein, J. R. Woolford, N. J. Lehrbach et al., 2008 Piwi and piRNAs act upstream of an endogenous siRNA pathway to suppress Tc*3* transposon mobility in the *Caenorhabditis elegans* germline. Mol. Cell 31: 79–90. https://doi.org/10.1016/j.molcel.2008.06.003

Denver, D. R., K. Morris, and W. K. Thomas, 2003 Phylogenetics in *Caenorhabditis elegans*: an analysis of divergence and outcrossing. Mol. Biol. Evol. 20: 393–400. https://doi.org/10.1093/molbev/msg044

Dey, A., C. K. W. Chan, C. G. Thomas, and A. D. Cutter, 2013 Molecular hyperdiversity defines populations of the nematode *Caenorhabditis brenneri*. Proc. Natl. Acad. Sci. USA 110: 11056–11060. https://doi.org/10.1073/pnas.1303057110

Dolgin, E. S., and B. Charlesworth, 2008 The effects of recombination rate on the distribution and abundance of transposable elements. Genetics 178: 2169–2177. https://doi.org/10.1534/genetics.107.082743

Dubie, J. J., A. R. Caraway, M. K. M. Stout, V. Katju, and U. Bergthorsson, 2020 The conflict within: origin, proliferation and persistence of a spontaneously arising selfish mitochondrial genome. Phil. Trans. Royal Soc. B 375: 20190174. https://doi.org/10.1098/rstb.2019.0174

Duret, L., G. Marais, and C. Biémont, 2000 Transposons but not retrotransposons are located preferentially in regions of high recombination rate in *Caenorhabditis elegans*. Genetics 156:1661–1669. https://doi.org/10.1093/genetics/156.4.1661

Egilmez, N. K., R. H. Ebert 2nd, and R. J. Shmookler Reis, 1995 Strain evolution in Caenorhabditis elegans: transposable elements as markers of interstrain evolutionary history. J. Mol. Evol. 40: 372–381. https://doi.org/10.1007/BF00164023

Eide, D., and P. Anderson, 1985 Transposition of Tc*1* in the nematode *Caenorhabditis elegans*. Proc. Natl. Acad. Sci. USA 82: 1756–1760. https://doi.org/10.1073/pnas.82.6.1756

Eide, D., and P. Anderson, 1988 Insertion and excision of *Caenorhabditis elegans* transposable element Tc*1*. Mol. Cell. Biol. 8: 737–746. https://doi.org/10.1128/mcb.8.2.737-746.1988

Emmons, S. W., M. R. Klass, and D. Hirsh, 1979 Analysis of the constancy of DNA sequences during development and evolution of the nematode *Caenorhabditis elegans*. Proc. Natl. Acad. Sci. USA 76: 1333–1337. https://doi.org/10.1073/pnas.76.3.1333

Emmons, S. W., S. Roberts, and K. S. Ruan, 1986 Evidence in a nematode for regulation of transposon excision by tissue-specific factors. Mol. Gen. Genet. 202: 410–415. https://doi.org/10.1007/BF00333270.

Emmons, S. W., and L. Yesner, 1984 High-frequency excision of transposable element Tc*1* in the nematode *Caenorhabditis elegans* is limited to somatic cells. Cell 36: 599–605. https://doi.org/10.1016/0092-8674(84)90339-8

Emmons, S. W., L. Yesner, K. S. Ruan, and D. Katzenberg, 1983 Evidence for a transposon in *Caenorhabditis elegans*. Cell 32: 55–65. https://doi.org/10.1016/0092-8674(83)90496-8

Fatt, H. V., and E. C. Dougherty, 1963 Genetic control of differential heat tolerance in two strains of the nematode *Caenorhabditis elegans*. Science 141: 266–267. https://doi.org/10.1126/science.141.3577.266

Files, J. G., S. Carr, and D. Hirsh, 1983 Actin gene family of *Caenorhabditis elegans*. J. Mol. Biol. 164: 355–375. https://doi.org/10.1016/0022-2836(83)90056-6

Fischer, S. E. J., E. Wienholds, and R. H. A. Plasterk, 2003 Continuous exchange of sequence information between dispersed Tc*1* transposons in the *Caenorhabditis elegans* genome. Genetics 164: 127–134. https://doi.org/10.1093/genetics/164.1.127

Guindon, S., J. F. Dufayard, V. Lefort, M. Anisimova, W. Hordijk et al., 2010 New algorithms and methods to estimate maximum-likelihood phylogenies: assessing the performance of PhyML 3.0. Syst. Biol. 59: 307−321. https://doi.org/10.1093/sysbio/syq010

Han, M., G. Wei, C. E. McManus, L. W. Hillier, and V. Reinke, 2019 Isolated *C. elegans* germ nuclei exhibit distinct genomic profiles of histone modification and gene expression. BMC Genomics 20:500. https://doi.org/10.1186/s12864-019-5893-9

Han, S., P. J. Basting, G. B. Dias, A. Luhur, A. C. Zelhof et al., 2021 Transposable element profiles reveal cell line identity and loss of heterozygosity in Drosophila cell culture. Genetics 219: iyab113. https://doi.org/10.1093/genetics/iyab113

Hill, W. G., and A. Robertson, 1966 The effect of linkage on limits to artificial selection. Genet. Res. 8: 269−294.

Hodgkin, J., and T. Doniach, 1997 Natural variation and copulatory plug formation in *Caenorhabditis elegans*. Genetics 146: 149–164. https://doi.org/10.1093/genetics/146.1.149

Holm, S., 1979 A simple sequentially rejective multiple test procedure. Scand. J. Stat. 6: 65–70.

Houle, D., and S. V. Nuzhdin, 2004 Mutation accumulation and the effect of copia insertions in Drosophila melanogaster. Genet. Res. 83: 7–18. https://doi.org/10.1017/s0016672303006505

Katju, V., L. B. Packard, L. Bu, P. D. Keightley, and U. Bergthorsson, 2015 Fitness decline in spontaneous mutation accumulation lines of *Caenorhabditis elegans* with varying effective population sizes. Evolution 69: 104−116. https://doi.org/10.1111/evo.12554

Katju, V., L. B. Packard, and P. D. Keightley, 2018 Fitness decline under osmotic stress in *Caenorhabditis elegans* populations subjected to spontaneous mutation accumulation at varying population sizes. Evolution 72: 1000−1008. https://doi.org/10.1111/evo.13463

Keane, T. M., K. Wong, and D. J. Adams, 2013 RetroSeq: transposable element discovery from next-generation sequencing data. Bioinformatics 29: 389−390. https://doi.org/10.1093/bioinformatics/bts697

Kidwell, M. G., and J. B. Novy, 1979 Hybrid dysgenesis in *Drosophila melanogaster*: sterility resulting from gonadal dysgenesis in the P-M system. Genetics 92: 1127−1140. https://doi.org/10.1093/genetics/92.4.1127

Klein, S. J., and R. J. O’Neill, 2018 Transposable elements: genome innovation, chromosome diversity, and centromere conflict. Chromosome Res. 26: 5–23. https://doi.org/10.1007/s10577-017-9569-5

Konrad, A., S. Flibotte, J. Taylor, R. H. Waterston, D. G. Moerman et al., 2018 Mutational and transcriptional landscape of spontaneous gene duplications and deletions in *Caenorhabditis elegans*. Proc. Natl. Acad. Sci. USA 115: 7386−7391. https://doi.org/10.1073/pnas.1801930115

Konrad, A., O. Thompson, R. H. Waterston, D. G. Moerman, P. D. Keightley et al., 2017 Mitochondrial mutation rate, spectrum and heteroplasmy in *Caenorhabditis elegans* spontaneous mutation accumulation lines of differing population size. Mol. Biol. Evol. 34: 1319–1334. https://doi.org/10.1093/molbev/msx051

Korswagen, H. C., R. M. Durbin, M. T. Smits, and R. H. Plasterk, 1996 Transposon Tc*1*-derived, sequence-tagged sites in *Caenorhabditis elegans* as markers for gene mapping. Proc. Natl. Acad. Sci. USA 93: 14680–14685. https://doi.org/10.1073/pnas.93.25.14680

Langley, C. H., E. Montgomery, R. Hudson, N. Kaplan, and B. Charlesworth, 1988 On the role of unequal exchange in the containment of transposable element copy number. Genet. Res. 52: 223–235. https://doi.org/10.1017/s0016672300027695.

Laricchia, K. M., S. Zdraljevic, D. E. Cook, and E. C. Andersen, 2017 Natural variation in the distribution and abundance of transposable elements across the *Caenorhabditis elegans* species. Mol. Biol. Evol. 34: 2187–2202. https://doi.org/10.1093/molbev/msx155

Lee, H.-C., W. Gu, M. Shirayama, E. Youngman, D. Conte Jr. et al. 2012 *C. elegans* piRNAs mediate the genome-wide surveillance of germline transcripts. Cell 150: 78–87. https://doi.org/10.1016/j.cell.2012.06.016

Lee, Y., W. Hwang, J. Jung, S. Park, J. J. T. Cabatbat et al., 2016 Inverse correlation between longevity and developmental rate among wild *C. elegans* strains. Aging 8: 986–999. https://doi.org/10.18632/aging.100960

Levitt, A., and S. W. Emmons, 1989 The Tc2 transposon in Caenorhabditis elegans. Proc. Natl. Acad. Sci. USA 86: 3232−3236. https://doi.org/10.1073/pnas.86.9.3232

Liao, L. W., B. Rosenzweig, and D. Hirsh, 1983 Analysis of a transposable element in *Caenorhabditis elegans*. Proc. Natl. Acad. Sci. USA 80: 3585−3589. https://doi.org/10.1073/pnas.80.12.3585

Lipinski, K. J., J. C. Farslow, K. A. Fitzpatrick, M. Lynch, V. Katju et al., 2011 High spontaneous rate of gene duplication in *Caenorhabditis elegans*. Curr. Biol. 21: 306−310. https://doi.org/10.1016/j.cub.2011.01.026

Liu, T., A. Rechtsteiner, T. A. Egelhofer, A. Vielle, I. Latorre et al., 2011 Broad chromosomal domains of histone modification patterns in C. elegans. Genome Res. 21: 227−236. https://doi.org/10.1101/gr.115519.110

Lynch, M., 1985 Spontaneous mutations for life-history characters in an obligate parthenogen. Evolution 39: 804–818. https://doi.org/10.1111/j.1558-5646.1985.tb00422.x

Malone, C. J., L. Misner, N. Le Bot, M.-C. Tsai, J. M. Campbell et al., 2003 The *C. elegans* hook protein, ZYG-12, mediates the essential attachment between the centrosome and nucleus. Cell 115: 825−836. https://doi.org/10.1016/s0092-8674(03)00985-1

McClintock, B., 1950 The origin and behavior of mutable loci in maize. Proc. Natl. Acad. Sci. USA 36: 344–355. https://doi.org/10.1073/pnas.36.6.344

McMurchy, A. N., P. Stempor, T. Gaarenstroom, B. Wysolmerski, Y. Dong et al., 2017 A team of heterochromatin factors collaborates with small RNA pathways to combat repetitive elements and germline stress. Elife. 6: e21666. https://doi.org/10.7554/eLife.21666

Moerman, D. G., G. M. Benian, R. J. Barstead, L. A. Schriefer, and R. H. Waterston, 1988 Identification and intracellular localization of the unc-22 gene product of Caenorhabditis elegans. Genes Dev. 2:93−105. https://doi.org/10.1101/gad.2.1.93

Moerman, D. G., G. M. Benian, and R. H. Waterston, 1986 Molecular cloning of the muscle gene unc-22 in Caenorhabditis elegans by Tc1 transposon tagging. Proc. Natl. Acad. Sci. USA 83: 2579−2583. https://doi.org/10.1073/pnas.83.8.2579

Moerman, D. G., and R. H. Waterston, 1984 Spontaneous unstable *unc-22* IV mutations in *C. elegans* var. Bergerac. Genetics 108: 859–877. https://doi.org/10.1093/genetics/108.4.859

Mori, I., D. G. Moerman, and R. H. Waterston, 1988 Analysis of a mutator activity necessary for germline transposition and excision of Tc1 transposable elements in Caenorhabditis elegans. Genetics 120: 397−407. https://doi.org/10.1093/genetics/120.2.397

Muñoz-López, M., and J. L. García-Pérez, 2010 DNA transposons: nature and applications in genomics. Curr. Genomics 11: 115−128. https://doi.org/10.2174/138920210790886871

Nelson, M. G., R. S. Linheiro, and C. M. Bergman, 2017 McClintock: an integrated pipeline for detecting transposable element insertions in whole-genome shotgun sequencing data. G3 (Bethesda) 7: 2763. https://doi.org/10.1534/g3.117.043893

Nicholas, W. L., E. C. Dougherty, and E. L. Hansen, 1959 Axenic cultivation of *C. briggsae* (Nematoda: Rhabditidae) with chemically undefined supplements; comparative studies with related nematodes. Ann. N.Y. Acad. Sci. 77: 218–236. https://doi.org/10.1111/j.1749-6632.1959.tb36902.x

Nigon, V., 1949 Les modalités de la reproduction et le déterminisme du sexe chez quelques nematodes libres. Ann. Sci. Nat. Zool. Biol. Anim. 11: 1–132.

Nigon, V., and E. C. Dougherty, 1949 Reproductive patterns and attempts at reciprocal crossing of Rhabditis elegans Maupas, 1900, and Rhabditis briggsae Dougherty and Nigon, 1949 (Nematoda: Rhabditidae). J. Exp. Zool. 112: 485–503. https://doi.org/10.1002/jez.1401120307

Nigon, V. M., and M.-A. Félix, 2017 History of research on *C. elegans* and other free-living nematodes as model organisms. WormBook 2017: 1–84. https://doi.org/10.1895/wormbook.1.181.1

Orgel, L. E., and F. H. Crick, 1980 Selfish DNA: the ultimate parasite. Nature 284: 604−607. https://doi.org/10.1038/284604a0

Pasyukova, E. G., S. V. Nuzhdin, T. V. Morozova, and T. F. C. Mackay, 2004 Accumulation of transposable elements in the genome of *Drosophila melanogaster* is associated with a decrease in fitness. *J*. Heredity 95: 284–290. https://doi.org/10.1093/jhered/esh050 Payer

L. M., and K. H. Burns, 2019 Transposable elements in human genetic disease. Nat. Rev. Genet. 20: 760−772. https://doi.org/10.1038/s41576-019-0165-8

Plasterk, R. H., Z. Izsvák, and Z. Ivics, 1999 Resident aliens: the Tc*1*/*mariner* superfamily of transposable elements. Trends Genet. 15: 326−332. https://doi.org/10.1016/s0168-9525(99)01777-1

Quinlan, A. R., and I. M. Hall, 2010 BEDTools: a flexible suite of utilities for comparing genomic features. Bioinformatics 26: 841–842. https://doi.org/10.1093/bioinformatics/btq033

R Core Team, 2014. R: A language and environment for statistical computing. R Foundation for Statistical Computing, Vienna, Austria. URL http://www.R-project.org/.

Rebollo, R., S. Farivar, and D. L. Mager, 2012 C-GATE — catalogue of genes affected by Transposable elements. Mobile DNA 3: 9. https://doi.org/10.1186/1759-8753-3-9

Reed, K. J., J. M. Svendsen, K. C. Brown, B. E. Montgomery, T. N. Marks et al., 2020 Widespread roles for piRNAs and WAGO-class siRNAs in shaping the germline transcriptome of *Caenorhabditis elegans*. Nucl. Acids Res. 48: 1811−1827. https://doi.org/10.1093/nar/gkz1178

Riddle, D. L., T. Blumenthal, B. J. Meyer, and J. R. Priess, editors, 1997 C. elegans II. 2nd edition. Cold Spring Harbor (NY): Cold Spring Harbor Laboratory Press; 1997. Section II, Origins of the Model. Available from: https://www.ncbi.nlm.nih.gov/books/NBK20127/

Rizzon, C., G. Marais, M. Gouy, and C. Biémont, 2002 Recombination rate and the distribution of transposable elements in the *Drosophila melanogaster* genome. Genome Res. 12: 400−407. https://doi.org/10.1101/gr.210802

Robb, S. M. C., L. Lu, E. Valencia, J. M. Burnette 3rd, Y. Okumoto et al., 2013 The use of RelocaTE and unassembled short reads to produce high-resolution snapshots of transposable element generated diversity in rice. G3 (Bethesda) 3: 949−957. https://doi.org/10.1534/g3.112.005348

Rockman, M. V., and L. Kruglyak, 2009 Recombinational landscape and population genomics of Caenorhabditis elegans. PLoS Genet. 5: e1000419. https://doi.org/10.1371/journal.pgen.1000419

Rosenzweig, B., L. W. Liao, and D. Hirsh, 1983a Sequence of the *C. elegans* transposable element Tc*1*. Nucleic Acids Res. 11: 4201–4209. https://doi.org/10.1093/nar/11.12.4201

Rosenzweig, B., L. W. Liao, and D. Hirsh, 1983b. Target sequences for the *C. elegans* transposable element Tc*1*. Nucleic Acids Res. 11: 7137–7140. https://doi.org/10.1093/nar/11.20.7137

Shook, D. R., and T. E. Johnson, 1999 Quantitative trait loci affecting survival and fertility-related traits in *Caenorhabditis elegans* show genotype-environment interactions, pleiotropy and epistasis. Genetics 153: 1233–1243. https://doi.org/10.1093/genetics/153.3.1233

Sijen, T., and R. H. Plasterk, 2003 Transposon silencing in the *Caenorhabditis* elegans germ line by natural RNAi. Nature 426: 310–314. https://doi.org/10.1038/nature02107

Sivasundar, A., and J. Hey, 2003 Population genetics of *Caenorhabditis elegans*: the paradox of low polymorphism in a widespread species. Genetics 163:147–157. https://doi.org/10.1093/genetics/163.1.147

Sokal, R. R., and F. J. Rohlf, 1995 *Biometry: The Principles and Practices of Statistic in Biological Research*. W. H. Freeman and Company, New York.

Sulston, J. E., and S. Brenner, 1974 The DNA of *Caenorhabditis elegans*. Genetics 77: 95−104. https://doi.org/10.1093/genetics/77.1.95

Sultana, T., A. Zamborlini, G. Cristofari, and P. Lesage, 2017 Integration site selection by retroviruses and transposable elements in eukaryotes. Nat. Rev. Genet. 18: 292−308. https://doi.org/10.1038/nrg.2017.7

Thomas, C. G., W. Wang, R. Jovelin, R. Ghosh, T. Lomasko et al., 2015 Full-genome evolutionary histories of selfing, splitting, and selection in *Caenorhabditis*. Genome Res. 25: 667−678. https://doi.org/10.1101/gr.187237.114

van’t Hof, A. E., N. Edmonds, M. Dalíková, F. Marec, and I. J. Saccheri, 2011 Industrial melanism in British peppered moths has a singular and recent mutational origin. Science 20: 958–960. https://doi.org/10.1126/science.1203043

Vaser, R., S. Adusumalli, S. N. Leng, M. Sikic, and P. C. Ng, 2016 SIFT missense predictions for genomes. Nat. Protoc. 11: 1–9. https://doi.org/10.1073/10.1038/nprot.2015.123

Vastenhouw, N. L., S. E. J. Fischer, V. J. P. Robert, K. L. Thijssen, A. G. Fraser et al., 2003 A genome-wide screen identifies 27 genes involved in transposon silencing in *C. elegans*. Curr. Biol. 13: 1311–1316. https://doi.org/10.1016/s0960-9822(03)00539-6

Vendrell-Mir, P., F. Barteri, M. Merenciano, J. González, J. M. Casacuberta et al., 2019 A benchmark of transposon insertion detection tools using real data. Mob. DNA 10: 53. https://doi.org/10.1186/s13100-019-0197-9

Vertino, A., S. Ayyadevara, J. J. Thaden, and R. J. Shmookler Reis, 2011 A narrow quantitative trait locus in *C. elegans* coordinately affects longevity, thermotolerance, and resistance to paraquat. Front. Genet. 2: 63. https://doi.org/10.3389/fgene.2011.00063

Wallis, D. C., D. A. H. Nguyen, C. J. Uebel, and C. M. Phillips, 2019 Visualization and quantification of transposon activity in *Caenorhabditis elegans* RNAi pathway mutants. G3 (Bethesda) 9: 3825–3832. https://doi.org/10.1534/g3.119.400639

Weick, E.-M., and E. A. Miska, 2014 piRNAs: from biogenesis to function. Development 141: 3458–3471. https://doi.org/10.1242/dev.094037

Wells, J. N., and C. Feschotte, 2020 A field guide to eukaryotic transposable elements. Annu. Rev. Genet. 54: 539–561. https://doi.org/10.1146/annurev-genet-040620-022145

Wood, W. B., R. Hecht, S. Carr, R. Vanderslice, N. Wolf et al., 1980 Parental effects and phenotypic characterization of mutations that affect early development in *Caenorhabditis elegans*. Dev. Biol. 74: 446–469. https://doi.org/10.1016/0012-1606(80)90445-5

Yigit, E., P. J. Batista, Y. Bei, K. M. Pang, C. C. Chen et al., 2006 Analysis of the *C. elegans* Argonaute family reveals that distinct argonautes act sequentially during RNAi. Cell 127: 747–757. https://doi.org/10.1016/j.cell.2006.09.033

Yu, T., X. Huang, S. Dou, X. Tang, S. Luo et al., 2021 A benchmark and an algorithm for detecting germline transposon insertions and measuring de novo transposon insertion frequencies. Nucleic Acids Res. 49: e44. https://doi.org/10.1093/nar/gkab010

Yuhang, Z., T. C. Cheng, G. Huang, Q. Lu, M. D. Surleac et al., 2019 Transposon molecular domestication and the evolution of the RAG recombinase. Nature 569: 79–84. https://doi.org/10.1038/s41586-019-1093-7

